# Enhancing Sox/Oct cooperativity induces higher-grade developmental reset

**DOI:** 10.1101/2022.09.23.509242

**Authors:** Caitlin M. MacCarthy, Vikas Malik, Guangming Wu, Taras Velychko, Gal Keshet, Ralf Jauch, Vlad Cojocaru, Hans R. Schöler, Sergiy Velychko

**Affiliations:** Max Planck Institute for Molecular Biomedicine, Münster, Germany; Department of Medicine, Columbia Center for Human Development, Columbia Stem Cell Initiative, Herbert Irving Comprehensive Cancer Center, Columbia University Irving Medical Center, New York, USA; Guangzhou Laboratory, Guangzhou International Bio Island, Guangzhou, China; Max Planck Institute for Biophysical Chemistry, Göttingen, Germany; Hebrew University of Jerusalem, Israel; School of Biomedical Sciences, Li Ka Shing Faculty of Medicine, The University of Hong Kong, China; University of Utrecht, The Netherlands; STAR-UBB Institute, Babeş-Bolyai University, Cluj-Napoca, Romania

## Abstract

The discovery of induced pluripotent stem cell (iPSC) technology by Shinya Yamanaka has truly enabled the stem cell field. After 16 years of intense research, the delivery methods and culture media have improved but the original factors—Oct4, Sox2, Klf4, and Myc (OSKM)—remain central for driving reprogramming.

Here we define structural elements in chimeric Sox2/Sox17 transcription factors that rescued the ability of nonfunctional Oct factors to induce pluripotency. Most importantly, we discovered a single amino acid swap in the DNA-binding domain of Sox2, A61V, that stabilizes the Sox/Oct heterodimer on DNA through hydrophobic interaction with Oct. The highly cooperative Sox2^AV^ mutant enables iPSC generation with Oct4 orthologs, such as Oct2 and Oct6, as well as rescues otherwise detrimental Oct4 mutants and domain deletions. Sox2^AV^ has a dramatic effect on the cell fate reset, significantly improving the developmental potential of OSKM iPSCs. Moreover, by swapping multiple beneficial elements of Sox17 into Sox2 we have built a chimeric super-SOX factor—Sox2-17—that delivers unprecedented reprogramming efficiency and kinetics in five tested species. Sox2-17 enhances five-, four-, and three-factor reprogramming up to hundreds of times, enables two-factor generation of human iPSCs, and allows integration-free reprogramming of otherwise non-permissive aged human, non-human primate, and cattle fibroblasts.

Our study demonstrates that a complete developmental reset requires both robust activation of regulatory elements controlled by the canonical *SoxOct* motif and limiting cellular proliferation driven by Oct4 and Myc. A high level of Sox2 expression and Sox2/Oct4 heterodimerization emerge as the key determinants of high-grade pluripotency that fades along the naïve-to-primed continuum. Transient expression of SK cocktail can restore the naivety, providing a powerful technology to induce more complete developmental reset in pluripotent cells across species.

## INTRODUCTION

The ability of some transcription factors (TFs), such as MyoD, to convert cell fates has been known since 1987^1^. It was not until 2006 that the discovery of induced pluripotent stem cells (iPSCs) by Takahashi and Yamanaka^2^ brought TF-based cell fate conversion into the spotlight, and rightly so. Pluripotent stem cells (PSCs) are unique in their ability to give rise to all the tissues of the animal body. They are the most primordial cell type of our bodies, the “stemmiest” stem cells. The induction of pluripotency amounts to the ultimate cell rejuvenation, once thought to be impossible^3^. iPSC technology has already made enormous contributions to human developmental studies, allowed new strategies for drug discovery, and even provided a source for cell replacement therapy. However, the most radical endeavor is perhaps yet to come: recent studies demonstrated the ability of reprogramming factors to reverse aging on the whole-organism level in mice^4–9^; time-restricted reprogramming factor induction could also rejuvenate human cells^10^.

Oct4, Sox2, Klf4, and cMyc (OSKM) - all components of the Yamanaka cocktail, control the pluripotency of the inner cell mass of the blastocyst^11,12^. We learned how to artificially expand PSCs from the inner cell mass in the form of embryonic stem cells (ESCs)^13^. In natural development, this stage is very transient, the factors we use for reprogramming evolved to reset the inner cell mass of the embryo, but simultaneously allow or even drive the differentiation. Oct4, Sox2, and Klf4 (OSK) are pioneer TFs capable of opening silent chromatin; iPSC technology harnesses their pioneering ability to reprogram cell fate^14,15^. Oct4 stands out as the master regulator of the pluripotency network. Knock out of Oct4 in ESCs leads to an inevitable collapse of pluripotency^16,17^; and forced expression of Oct4 can partially compensate for the loss of Sox2, albeit functional compensation by other Sox factors could not be excluded^18^. Oct4 was considered to be the only factor that cannot be replaced by other members of its family in iPSC generation^19,20^. Endogenous Oct4 activation occurs relatively early in reprogramming process, and it is endogenous Sox2 activation that signifies the completion of pluripotency induction^21^. Moreover, we showed that expression of exogenous Oct4 during reprogramming leads to worse developmental potential of OSKM versus SKM iPSCs^22^. Oct4 plays divergent roles in establishing pluripotency during mouse and human development: Oct4 knockout mouse blastocysts still develop a Nanog^+^ inner cell mass, while human OCT4-null blastocysts fail to do so^23,24^. Correspondingly, SKM induction is sufficient to induce pluripotency in mouse somatic cells^22,25^. However, SKM reprogramming has not been demonstrated for human or other species, emphasizing the need to develop alternative strategies to improve non-murine reprogramming.

Oct4 cooperates with Sox2 to co-regulate the majority of its targets in pluripotent cells^26^. Sox2/Oct4 cooperativity is mediated by protein-protein interaction between their DNA-binding domains and by DNA allostery^27^. At the beginning of reprogramming to iPSCs, when native Oct4 and Sox2 sites are inaccessible, each factor often binds independently^15^, yet the sites engaged by both factors are most likely to be opened^28,29^. Sox2/Oct4 cooperativity, particularly on *HoxB1*-like canonical *SoxOct* motifs, was shown to be essential for the induction and maintenance of pluripotency^30^. On the other hand, Sox17 cooperates with Oct4 on compressed, non-canonical *SoxOct* motif and cannot induce pluripotency but rather controls primitive endoderm and germline specification^19,31–36^. Jauch et al. discovered that a single residue swap between Sox17 and Sox2, glutamate to lysine at HMG box position 57 (Sox17^E57K^), shifts its binding preference to the canonical *SoxOct* motif converting Sox17 into a pluripotency inducer^36^. The Sox17 C-terminus transactivator (CTD) is larger and more potent than Sox2-CTD. The Sox17-CTD alone can enhance the reprogramming ability of Sox2^31^. Here we found that replacing Sox2 with the Sox17^E57K^ in the reprogramming cocktail rescues otherwise detrimental Oct4 mutants and allows reprogramming with POU factors other than Oct4. We generated a library of chimeric Sox2-Sox17 TFs to find the structural elements of Sox17 responsible for this peculiar phenotype. The project led us to a new understanding of reprogramming factors’ structure/function paradigm and allowed us to build enhanced cell fate reprogramming machinery that does not occur in nature.

## RESULTS

### Defining structural elements of Sox17 that benefit induction of pluripotency

Oct4 (Pou5f1) is the only TF of the POU (Pit1, Oct1/Oct2, UNC-86) family that can induce pluripotency in mouse and human cells^2,37,38^, while other members of the family, such as the ubiquitously expressed Oct1 (Pou2f1), neural Oct6 (Pou3f1), and Brn4 (Pou3f3 or Oct9) cannot^19,20,39^. POU factors have different binding profiles and different preferences for hetero-versus homodimerization^20,28,40^. In our search to find what makes Oct4 unique among POU factors, we discovered that the Sox17^E57K^ (Sox17^EK^) mutant^36^, but not wild-type Sox2, can efficiently generate iPSCs in combination with Brn4 (Fig. 1a). POU factors consist of a DNA-binding domain (POU domain), flanked by N- and C-terminal transactivators (NTD and CTD). The POU domain is bipartite, consisting of a POU-specific domain (POU_S_) and a POU-homeodomain (POU_HD_) joined by a flexible non-conserved linker. The Oct4 but not the Oct1 linker contains an alpha-helix near its N-terminus^41,42^. Replacement of the Oct4 linker with those from other POU factors, or even the point mutant L80A in the linker helix is detrimental for induction and maintenance of pluripotency^41,43,44^. Surprisingly, Sox17^EK^ could also rescue the reprogramming ability of Oct4^L80A^ (Fig. S1a-b).

**Fig. 1.**
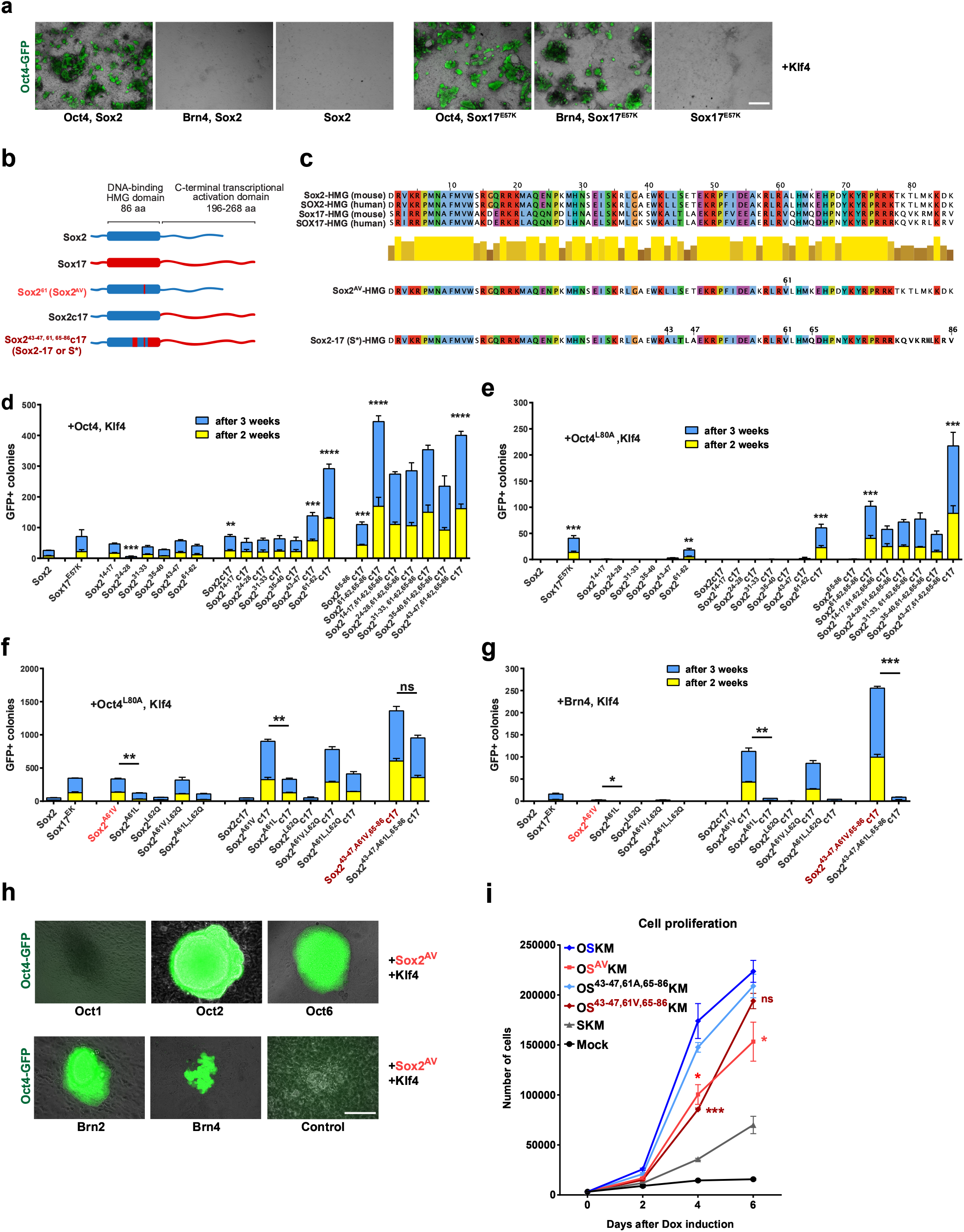
Reprogramming screen of Sox2-Sox17 chimeric TF library. **a**, Brightfield and Oct4-GFP merged overview images showing retroviral reprogramming of MEFs carrying Oct4-GFP (OG2) reporter on 21 dpi (scale=l mm). **b**, Schematic representation of Sox2 and Sox17 structures and chimeric transcription factors (TFs) generated by swapping nonconserved residues from Sox17 into Sox2. Sequence from Sox2 in blue and from Sox17 in red. **c**, Protein sequence alignment of DNA binding domains of mouse and human Sox2, Sox17 as well as the most crucial chimeric Sox factors of this study. **d-g**, Reprogramming of Oct4-GFP MEFs by retroviral vectors carrying Klf4, Sox2-Sox17 chimeric TFs, and wild-type Oct4 (**d**), Oct4^L80A^ linker mutant (**e-f**), or Brn4 (**g**). Error bars represent SD; n = 3. Statistical significance was calculated with Student’s t-test. **h**, Representative phase-contrast and Oct4-GFP merged microscopy images of primary iPSCs colonies generated with retroviral vectors carrying different POU factors combined with Sox2^AV^ and Klf4 (Scale=200μm). **i**, Cell proliferation assay, where 2×10^3^ MEFs were transduced with the indicated tet-inducible polycistronic constructs in 96-well plates. Error bars represent SD; n = 3. The cells were counted after 2, 4, and 6 dpi. 61V was compared with 61A for each construct to calculate the statistical significance with Student’s t-test.

In order to find the structural elements responsible for this rescue, we constructed a library of chimeric Sox2-Sox17 TFs, where we swapped the non-conserved residues from Sox17 into Sox2 (Fig. 1b-c). Monocistronic retroviral supernatants were used for reprogramming mouse embryonic fibroblasts (MEFs) carrying endogenous Oct4-GFP reporter (OG2); the volumes of viral supernatants were adjusted based on qPCR titration (Fig. S1c-d). Initial screening results pointed to residues 61-62 of Sox17 as the most crucial (Fig. 1d). The Sox17-CTD enhanced reprogramming with wild-type Oct4, but could not rescue Oct4^L80A^ (Sox2c17 chimera, Fig. 1d-e) unless combined with the 61-62 region (Sox2^61–62^c17 chimera, Fig. 1d-e). Two other Sox17 regions, 43-47 and 65-86, also boosted the reprogramming efficiency, particularly when combined with 61-62 (Fig. 1d-e). Other Sox17 elements either did not affect reprogramming or, in case of residues 24-28, significantly decreased reprogramming efficiency (Fig. 1d). Further analysis determined that a single A61V swap in Sox2 rescued both Oct4^L80A^ and Brn4 reprogramming, while L62Q was inconsequential (Fig. 1f-g). Sox2^A61V^ also allowed reprogramming with Oct2 (Pou2f2), Oct6, and Brn2 (Pou3f2 or Oct7), leaving Oct1 (Pou2f1) as the only tested POU factor that did not yield iPSC colonies when co-expressed with Sox2^A61V^ and Klf4 in MEFs (Fig. 1h). We also tested the A61L mutation, as leucine has even higher hydrophobicity than valine. While Sox2^A61L^ performed better than wild-type Sox2, it was significantly worse than Sox2^A61V^ (from now on Sox2^AV^), especially in reprogramming with Brn4 (Fig. 1f-g). Notably, a complex chimeric Sox factor, Sox2^43–47,61,65–86^c17 (Sox2-17), that incorporates 16 beneficial Sox17 residues into the Sox2 HMG-box domain as well as a complete CTD swap (Fig. 1b-c) exceeded the reprogramming ability of Sox2, Sox2c17, or Sox17^EK^ (Fig. 1f-g). Cell proliferation is essential for the induction of pluripotency; cMyc, GATA factors, and the SV40 large T antigen increase reprogramming efficiency by boosting cell proliferation^22,45^. However, the A61V mutation has the opposite effect: four-factor induction with Sox2^AV^ or Sox2-17 resulted in significantly lower cell proliferation compared to their respective 61A variants (Fig. 1i). The repressive effect on cell proliferation explains why A61V, while being able to rescue nonfunctional Oct4 mutants, does not by itself increase reprogramming efficiency with wild-type Oct4 (Fig. 1d). The efficiency boost comes from the synergy between A61V and the stronger Sox17 transactivator (Fig. 1d-g).

### Sox2^A61V^ enhances Sox/POU cooperativity

The residue A61 is located in the third helix of the high-mobility group (HMG) domain of Sox2 that faces towards the POU_S_ domain when the heterodimer is bound to a canonical *SoxOct* motif (Fig. 2a-b). Compared to alanine, valine has two additional methyl groups, making it more hydrophobic. Molecular dynamic simulations (MDS) of the Sox/Oct heterodimer on the *HoxB1 SoxOct* DNA element showed that swapping A to V increases the average number of hydrophobic interactions between Sox2-HMG^61^ and Oct4- or Oct6-POU_S_ (Fig. 2c). The 121 residue of both Oct factors engaged with 61V the most (Fig. 2d), which could potentially increase the cooperativity of Sox2^AV^ with POU factors. The simulations also revealed a previously undescribed structural rearrangement of the Sox2/Oct4 heterodimer that lead to higher Sox2^61^/Oct4 ligancy. This rearrangement was not observed when Oct4 was replaced with Oct6 (Fig. 2c). In this configuration, stabilized by A61V, the residue 61 interacts with the core of the helix near G24 of Oct4-POU_S_ and residues R50 and K57 of HMG form salt bridges with E82 and E78 of the Oct4 linker (Fig. 2e). We termed this structure of Sox2/Oct4 heterodimer the SL configuration, as it involves both the POU_S_ and Linker of Oct4, as opposed to the S configuration, which involves only the POU_S_. The SL configuration also emerged in MDS of wild-type the Sox2/Oct4 heterodimer on *Nanog* promoter locus DNA confirming the *HoxB1* results (Fig. S2a-b). We also modeled another configuration described for Sox/Oct heterodimer on the rarer *Fgf4* motif, where Sox and Oct sites are separated by a three base-pair gap^46^. Sox2 and Oct4 cooperate more distantly on the *Fgf4* motif, forming a **D**istant **S** (DS) configuration that involves Sox2’s R75^30^, T78, and T80, but not A61 (Fig. 2f).

**Fig. 2.**
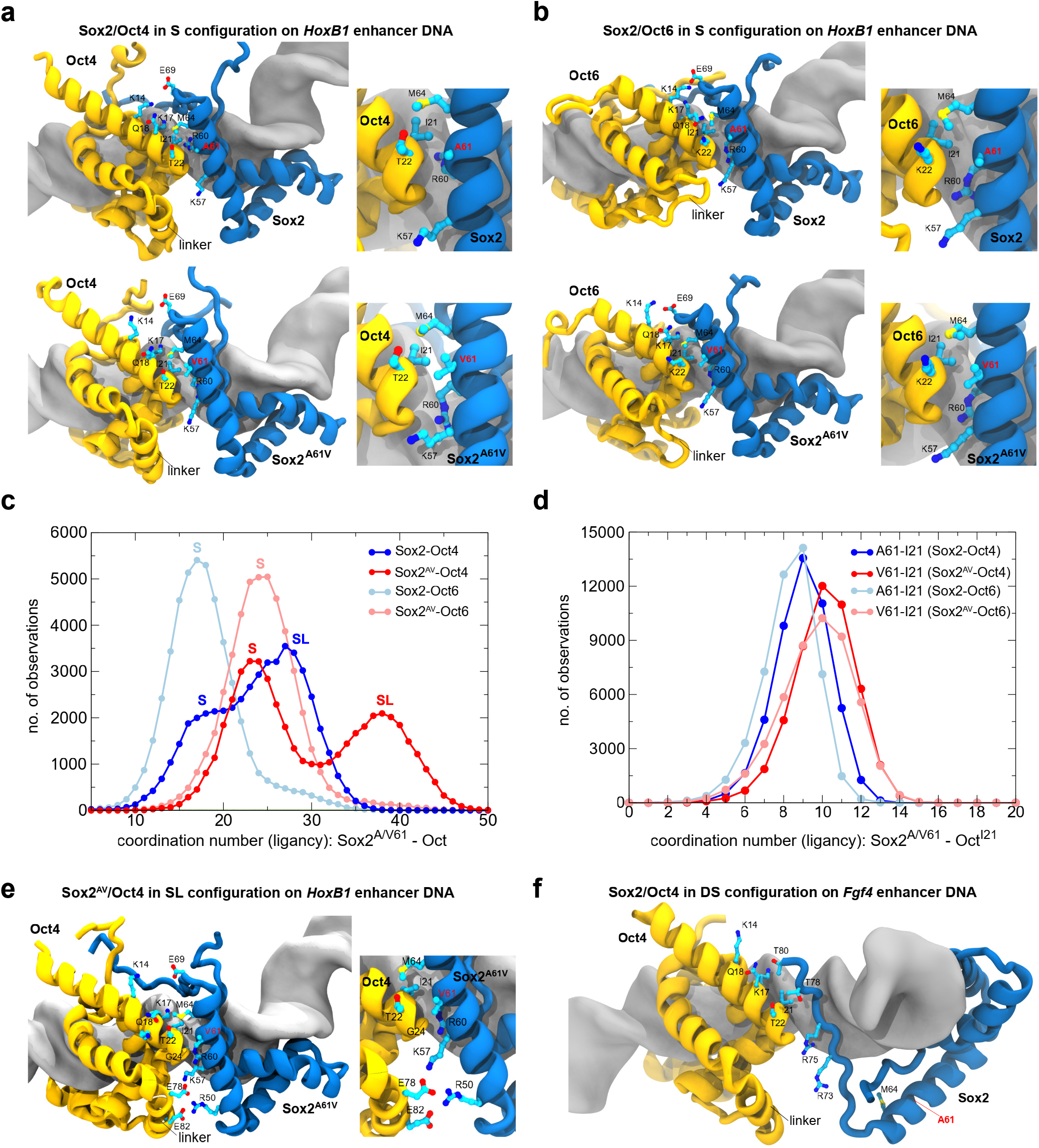
Molecular dynamic simulations reveals SL configuration of Sox/Oct4. **a-b**, A model of Sox2/Oct4 versus Sox2^AV^/Oct4 heterodimer (**a**) and Sox2/Oct6 versus Sox2^AV^/Oct6 heterodimer (**b**) in POU_S_ (S) configuration on *HoxB1* enhancer DNA containing canonical *SoxOct* motif. Only DNA-binding domains are shown. Oct4/Oct6 in yellow, Sox2/Sox2^AV^ in blue. **c-d**, Computer molecular dynamic simulations (MDS) of Sox/Oct heterodimers on *HoxB1* DNA. The plots show the coordination numbers (the number of contacts) between the residue 61 in Sox2 (blue) or Sox2^A61V^ (red) either with the entire DNA binding domain of Oct4 (dark) or Oct6 (light) molecule (**c**), or with residue 21 1 in Oct (**d**). Calculations were made for 4.8 μs of MDS of each ternary Sox/Oct/DNA complex. 4 independent 1.2 μs long simulations were performed using two different starting structural models (2 simulations per model). To ensure stochasticity, each simulation was started with a different distribution of atomic velocities. The distance threshold for a contact between two atoms was 4.5 A. **e**, A model of Sox2^AV^/Oct4 binding in POU_S_+Linker (SL) configuration, where V61 interacts with the helix core of POU_S_ at residue G24, and two salt bridges are formed between E78 and E82 of Oct4 linker and K57 and R50 of Sox2 HMG, respectively. **f**, A model of Sox2/Oct4 binding in **d**istant POU_S_ (DS) configuration on *Fgf4* motif; residue 61 of Sox2 is not involved.

We overexpressed FLAG-tagged Oct1, Oct2, Oct4, Oct6, Brn2, and Brn4 in HEK293 cells and after confirming their comparable expression (Fig. S3a) used the lysates for electromobility shift assays (EMSA) on the *Nanog* promoter locus containing a canonical *SoxOct* motif^47^. POU monomer binding was comparable for all tested factors with the exception of reduced binding by Oct1, but Oct4 showed the strongest heterodimerization capacity with Sox2 (Fig. S3b). There were no differences in monomer binding between wild-type Sox2, Sox2^AV^, and Sox2-17, but both Sox2^AV^ and Sox2-17 enhanced heterodimerization with all the tested POU factors (Fig. S3b). In order to find which domains of Oct4 remain vital for generating iPSCs when combined with Sox2^AV^ and other chimeric Sox factors, we first replaced all 17 residues of the Oct4 linker domain with synthetic poly-glycine linkers of different length (GL3-30) (Fig. 3a). Such flexible linkers were detrimental for reprogramming with wild-type Sox2, but Oct4^GL15-3^0 mutants were rescued by Sox2^AV^, and the efficiency was further increased by Sox2-17 (Fig. 3a). Second, we truncated either NTD and CTD of Oct4, both of which were previously shown to be crucial for reprogramming^48^. Indeed, neither Oct4ΔNTD nor Oct4ΔCTD could generate iPSCs when combined with wild-type Sox2 (Fig. 3b, S3c-d). However, Sox2^AV^ or Sox2c17 could rescue Oct4ΔCTD, and Sox2^AV^c17, Sox17^EK^, or Sox2-17 rescued both Oct4ΔNTD and Oct4ΔCTD (Fig. 3b, S3c-d). These results show that increased Sox/Oct cooperativity, especially when combined with a stronger Sox transactivator, can compensate for the loss of transactivation power by Oct4 in reprogramming. Third, none of the chimeric Sox factors could rescue the deletion of Oct4-POU_S_ domain, which is directly involved in Sox/Oct interaction (Fig. 3c). Remarkably, Sox2^AV^ could even give rise to a few GFP^+^ colonies with Oct4ΔPOU_HD_, where we truncated the POU_HD_ DNA-binding domain leaving the RKRKR peptide at its N-terminus that serves as nuclear localization signal (Fig. 3c, Fig. S3e). PCR-genotyping confirmed the identity of two clonal Oct4ΔPOU_HD_/Sox2^AV^/Klf4 iPSC lines (Fig. S3f) and they were able to contribute to chimeric mice (Fig. S3g) including the germline (Fig. S3h).

**Fig. 3.**
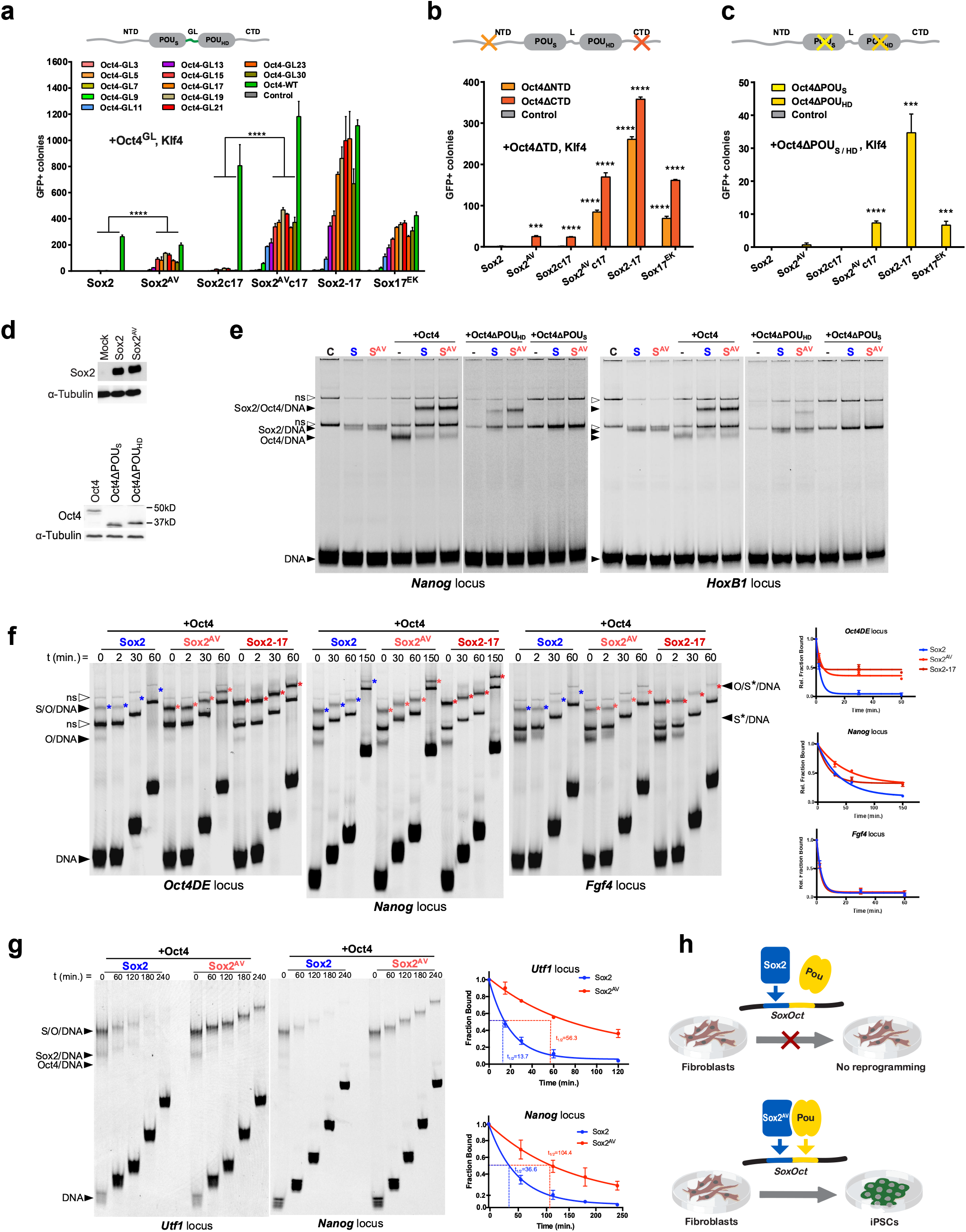
Enhanced Sox/Oct cooperativity rescues non-functional POU factors in reprogramming to pluripotency. **a-c**, OSK reprogramming of Oct4-GFP MEFs with monocistronic retroviral vectors carrying Oct4 domain deletion mutants where the linker domain was replaced with a synthetic poly-Glycine linker (GL) of different length (3-30 residues) (**a**), Oct4’s N- or C-terminal transactivator domain (NTD or CTD) was removed (**b**); POU_S_ or POU_HD_ (except for NLS) was removed (**c**) combined with wild-type Sox2 or Sox2-Sox17 chimeric factors. Error bars represent SD; n = 3. Statistical significance was calculated with Student’s t-test. **d**, Western blot of whole-cell lysates from HEK293 used in Fig. 3e. **e**, Electrophoretic mobility shift assays (EMSAs) of whole-cell lysates of HEK293 cells transfected with Sox2 (S, blue), Sox2^AV^ (S^AV^, light red) and wild-type, or mutant Oct4 in which the POU_S_ or POU_HD_ domain was removed on the *Nanog* promoter and *HoxB1* enhancer *SoxOct* DNA elements labeled with Cy5. White arrow heads indicate nonspecific bands (ns) and black arrow heads indicate free DNA or DNA bound by Oct4, Sox2, or the heterodimer. **f**, Representative kinetic off-rate EMSAs of whole-cell lysates from HEK293 overexpressing Oct4, Sox2 (blue), Sox2^A61V^ (light red) or Sox2^43–47,61,65–86^c17 (Sox2-17, S*, dark red) on *Oct4* distal enhancer (*Oct4DE), Nanog* promoter, or *Fgf4* enhancer DNA elements labeled with Cy5. White arrow heads indicated nonspecific bands (ns) and black arrow heads indicate free DNA or DNA bound by Oct4 (O/DNA), Sox (S/DNA), or Oct4/Sox heterodimer (O/S/DNA). Error bars on graphs represent SD; n = 3. **g**, Representative kinetic off-rate EMSAs using purified proteins bound to the *Utf1* enhancer and *Nanog* promoter loci labeled with Cy5. Following the binding reaction, halflife was determined by adding excess unlabeled *Nanog* element for the indicated time. The error bars on quantitation graphs represent SD, n =3. Ternary complex half-life t_1/2_. **l**, A schematic of the enabling effect of highly cooperative Sox2^AV^ mutant on reprogramming with tissue specific POU factors or incapable Oct4 mutants.

We overexpressed Oct4 and Sox2 mutants in HEK293 cells, confirmed equal expression (Fig. 3d), and performed whole-cell lysate EMSA experiments. Again, monomer binding showed no difference between Sox2^AV^ and wild-type Sox2 on either *HoxB1* or *Nanog* DNA elements (Fig. 3e), however, 61V increased the heterodimerization with wild-type Oct4 and partially rescued the DNA binding of the Oct4ΔPOU_HD_ mutant (Fig. 3e), in concordance with our reprogramming results (Fig 3c, Fig. S3e-h). The Oct4ΔPOU_HD_ rescue data are of particular interest because of previously debated reports on the role of POU subdomains in Oct4’s pioneering function. While a nucleosome does not pose a problem for Sox2 binding^49^, it does hinder canonical Oct4 binding because the POU_S_ and POU_HD_ engage opposite sides of DNA inevitably clashing with the histone core^47,50,51^. This has prompted speculations that Oct4 engages closed chromatin with just one of its domains^47^ or even that POU_S_ alone is involved in chromatin opening^51^. On the other hand, we showed that while Oct4 uses either the POU_S_ and POU_HD_ to recognize specific sequences on nucleosomes, the other domain scans unspecifically creating barriers to nucleosome closing^52^. The data presented here shows that removing POU_HD_ abolishes both Oct4 DNA binding on *SoxOct* motifs (Fig. 3e) and the reprogramming efficiency (Fig. 3c compared to 3a-b), highlighting the importance of POU_HD_ even in the context of highly-cooperative Sox factors. It appears unlikely that long-lived and, therefore, consequential Oct4 binding could be achieved with either POU_S_ or POU_HD_ alone in the context of wild-type Sox2 (Fig. 3e).

To compare the stability of Sox/Oct4 heterodimers on different natural regulatory *SoxOct* DNA elements, we performed off-rate EMSA experiments, where an excess of unlabeled DNA is added to pre-formed Sox/Oct/DNA complex and samples are loaded onto a gel over a time course. Both Sox2^AV^ and Sox2-17 dramatically enhanced the heterodimer stability on both Oct4 distal enhancer (*Oct4DE*) and *Nanog* promoter elements, but they showed similar to wild-type Sox2 stability on the *Fgf4* enhancer locus (Fig. 3f). Interestingly, a proportion of Sox2/Oct4/Oct4DE dissociated almost immediately, while the remaining complex was extraordinarily long-lived (Fig. 3f), suggesting the presence of at least two distinct Sox/Oct/DNA populations as with our MDS results (Fig. 2, S2). We used Oct4, Sox2, and Sox2^AV^ purified from insect cells (Fig. S3i) for EMSAs on three natural *SoxOct* elements: *Nanog* promoter^47^, *Utf1*^53^, and *Fgf4*^54^ enhancers (Fig. 3g, Fig. S3j). Sox2 and Sox2^AV^ monomer binding as well as heterodimerization on *Fgf4* were similar, while the heterodimerization on *Nanog* and *Utf1* were enhanced by the A61V mutation (Fig. S3j). The off-rate EMSAs on *Nanog* and *Utf1* elements are reminiscent of our whole-cell lysate experiments: A61V increased the stability of Sox2/Oct4/DNA complexes by 3 to 4 times (Fig. 3g). Sox2^AV^ also strongly increased otherwise diminished heterodimer stability with Oct4 linker mutants and Brn4 (Fig. S3k), explaining its rescue ability in reprogramming experiments (Fig. 1f-h, Fig. 3a). We conclude that POU factors and Oct4 mutants that are normally incapable of reprogramming to pluripotency can be rescued by forced cooperation with Sox2 (Fig. 3h).

Both Oct4 and Sox2 are pioneer factors capable of binding and activating gene expression within inaccessible chromatin^15,55^. We assembled a modified Widom 601 nucleosome sequence adapted with a *SoxOct* motif at the superhelical location (SHL) +6, which was used to resolve the Sox2/Oct4/nucleosome complex by cryo-electron microscopy^51^. Sox2^AV^ dramatically enhanced the stability of the Sox2/Oct4/nucleosome complex (Fig. 4a). We then performed chromatin immunoprecipitation with sequencing (ChlP-seq) to identify genome-wide binding sites at very early stage of reprogramming—two days after doxycycline (Dox)-induction of KS or OKS in MEFs. HOMER motif enrichment analysis^56^ showed that A61V samples were more enriched in *SoxOct* motif in both Oct4/Sox2^AV^/K and Oct6/Sox2^AV^/K Oct ChIP but there was no significant difference for Sox2 ChIP (Fig. S4a-b). Sox2 peaks were similar between Sox2 and Sox2^AV^ in both KS and OKS samples (Fig. 4b-c), suggesting that A61V does not change the binding profile of Sox2 itself; however, Oct4 binding was significantly enhanced in the presence of Sox2^AV^(Fig. 4b-c). ChIP for both Oct4 and Oct6 showed an increased number of peaks containing *SoxOct* motif in Sox2^AV^ compared to Sox2 samples (Fig. 4d), suggesting a redistribution in genome-wide binding of POU factors in the presence of Sox2^AV^. We performed TOBIAS footprinting analysis^57^ using publicly available assay for transposase-accessible chromatin using sequencing (ATAC-seq) dataset for ESCs versus MEFs^58^. Sox2/Oct4 footprints detected in ESC versus MEF ATAC-seq signal represent the genomic loci likely to be opened by the heterodimer during the reprogramming process. The occupancy Sox2^AV^ in those loci was only slightly stronger compared to Sox2, but Oct4 occupancy was dramatically increased (Fig. S4C). The stronger binding by Sox2^AV^/Oct4 compared to Sox2/Oct4 is illustrated by increased signal for both Oct4 and Sox2^AV^ occupancy at the key pluripotency targets such as *Gdf3, Nr5a2, Lefty2 Pou5f1*, and *Nanog* (Fig. 4e). Consistent with our modeling prediction (Fig. 2f) and EMSA results (Fig. 3f), binding at *Fgf4* locus was not affected (Fig. 4e).

**Fig. 4.**
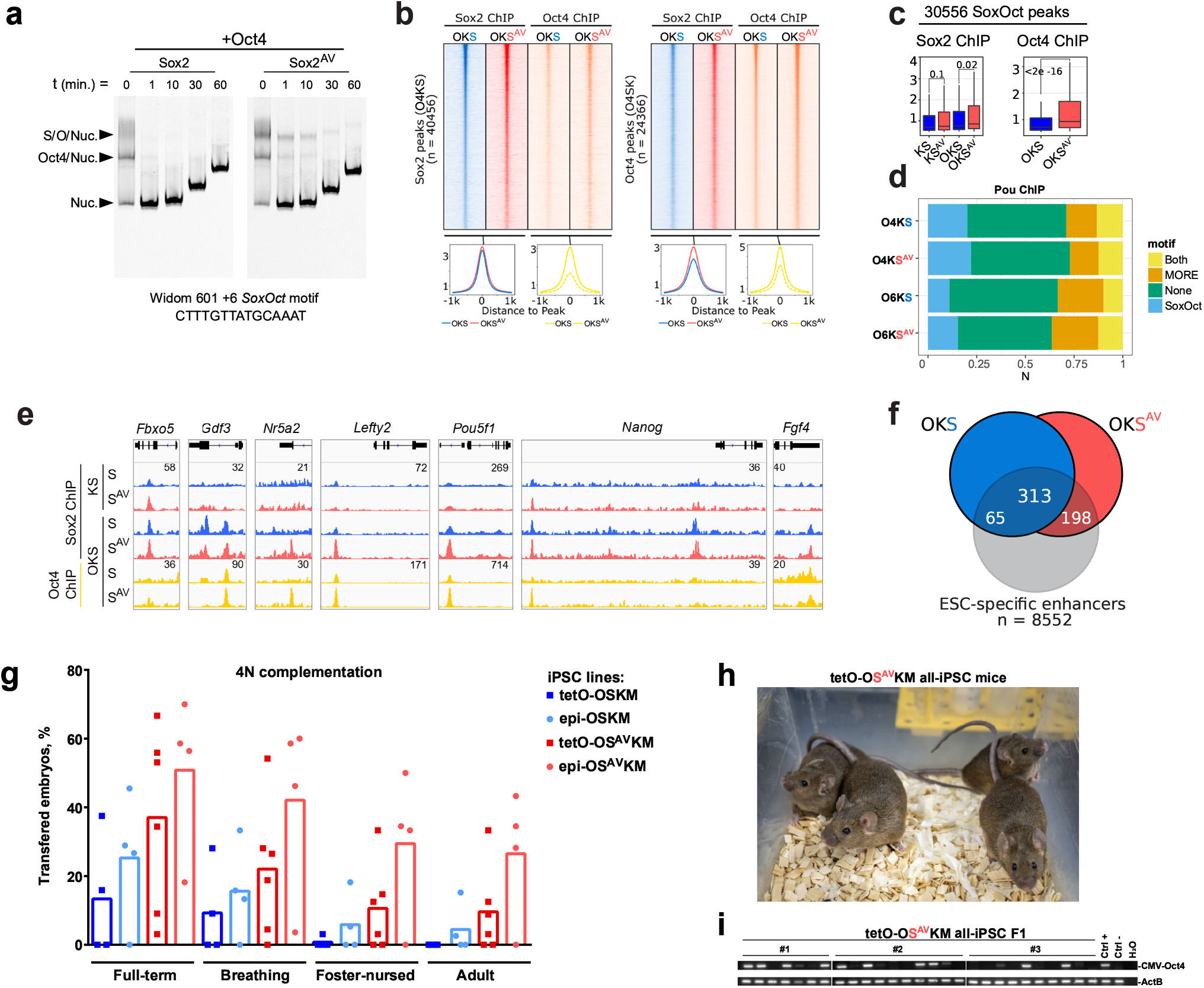
Highly-cooperative Sox2^AV^ improves the developmental potential of mouse OSKM iPSCs. **a**, Kinetic off-rate EMSAs of purified wild-type Sox2 or Sox2^AV^ in complex with Oct4 bound to the Widom 601 nucleosome sequence adapted with a *SoxOct* motif at superhelical location (SHL) +6. Following the binding reaction, ternary complex stability was determined by adding excess unlabeled *Nanog* element for the indicated time. Black arrow heads mark free nucleosome or the indicated nucleosome protein complex. **b**, Heat maps and read pileup plots of Sox2 and Oct4 ChlP-seq for 2 dpi tetO-OKS MEF reprogramming samples comparing Sox2^AV^ versus wild-type Sox2. **c**, Boxplots of RPM normalized ChlP-seq peaks for Oct4 and Sox2 for 2 dpi OKS and KS reprogramming samples. The midline indicates the median, boxes indicate the upper and lower quartiles and the whiskers indicate 1.5 times interquartile range. **d**, Fraction of binding sites containing *SoxOct, MORE*, both or none of the motifs between 2 dpi OKS reprogramming samples, where O is either Oct4 or Oct6, and S is either Sox2 or Sox2^AV^. **e**, Genome browser track of Oct4 and Sox2 ChlP-seq peaks for selected pluripotency-specific loci. **f**, Venn diagram showing a number of ESC-specific enhancers^65^ bound by Sox2 versus Sox2^AV^ on 2 dpi tetO-OKS reprogramming samples. **g**, Percentage of tetraploid (4N)-aggregated embryos derived from the indicated tet-inducible or integration-free episomal iPSCs that gave rise to full-term pups, pups that initiated breathing, pups that survived foster-nursing for at least 48h, and those survived to adulthood (at least 3 months). Bars represent the mean between all tested lines for each cocktail. **h**, Adult tetO-OS^AV^KM all-iPSC mice (9 months). **i**, PCR-genotyping of the progeny of tetO-OS^AV^KM all-iPSC mice.

In ESCs, Oct4 and Sox2 regulate genes cooperatively; the majority of pluripotency genes contain *SoxOct* motifs in their regulatory elements^59^. Oct4 also functions independently from Sox2 to drive ESC proliferation^60^. Overexpression of Oct4 in somatic cells induces hyperproliferation^61^. Accordingly, OSKM induction in MEFs caused a much stronger boost of cell proliferation compared to SKM induction (Fig. 1i)^22^. At the beginning of reprogramming, when Oct4 and Sox2 are overexpressed in somatic cells they tend to bind independently engaging thousands of nonnative genomic loci^15,28,29,62,63^. We hypothesized that enhancing Sox/Oct cooperativity could improve the reprogramming process, as cooperativity between TFs increases their specificity^64^. Indeed, already on day two of OKS induction, Sox2^AV^ engaged 511 of ESC-specific super-enhancers^65^, compared to 378 for wild-type Sox2 (Fig. 4f).

### Enhancing Sox2/Oct4 cooperativity improves the developmental potential of iPSCs

To assess the effect of enhanced Sox2/Oct4 cooperativity on the developmental potential of iPSCs, we performed tetraploid (4N)-complementation assays for iPSC lines generated using either lentiviral tet-inducible or episomal delivery methods, both polycistronic OSKM reprogramming cassettes carrying wild-type Sox2 or Sox2^AV^. Astonishingly, all 10 Sox2^AV^-generated iPSC lines could support full-term development (4N-on), giving rise to more than twice as many all-iPSC pups compared with wild-type Sox2 for both reprogramming methods (Fig. 4g, Table 1). The benefit of the highly-cooperative Sox2^AV^ was particularly evident for the survival of the all-iPSC mice. As we and others previously reported^22,66,67^, OSKM all-iPSC mice rarely survive to adulthood: none of the nine tetO-OSKM iPSC lines generated here or in our previous study gave rise to adult all-iPSC mice (Fig. 4g, Table 1, and^22^), while four out of six tetO-OS^AV^KM lines gave rise to adult all-iPSC mice with up to 33.3% efficiency (Fig. 4g-h, Table 1). The tetO-OS^AV^KM all-iPSC mice were fertile; PCR genotyping confirmed the inheritance of the transgene in their progeny (Fig. 4i). Episomal (epi-) vectors deliver milder overexpression, giving rise to better quality integration-free iPSCs, even in the presence of exogenous Oct4^22^. Yet, on average only 4.2% of transferred epi-OSKM embryos gave rise to adult all-iPSC mice, compared to 22.2% of epi-OS^AV^KM embryos. The top-quality integration-free OS^AV^KM iPSC line outperformed the top-quality OSKM line by almost three-fold: 43.3% and 15.2% of transferred embryos giving rise to adult all-iPSC mice, respectively (Fig. 4g, Table 1). Thus, enhancing Sox2/Oct4 cooperativity dramatically improves the developmental potential of mouse OSKM iPSCs.

**Table 1.** Tetraploid complementation results.

| iPSC line | Karyotype | Aggregates transferred | Full-Term pups | Breathing | Survived after 48h | Survived after 3 months |
| --- | --- | --- | --- | --- | --- | --- |
| tetO-OSKM #1 | 40, XY | 30 | 0 | 0 | 0 | 0 |
| tetO-OSKM #2 | 40, XY | 32 | 12 | 9 | 1 | 0 |
| tetO-OSKM #3 | 40, XY | 44 | 7 | 4 | 0 | 0 |
| tetO-OSKM #4 | 40, XY | 44 | 0 | 0 | 0 | 0 |
| tetO-OS<sup>AV</sup>KM #1 | 40, XY | 24 | 16 | 13 | 8 | 8 |
| tetO-OS<sup>AV</sup>KM #2 | 40, XY | 32 | 11 | 9 | 1 | 1 |
| tetO-OS<sup>AV</sup>KM #3 | 40, XY | 34 | 19 | 9 | 5 | 3 |
| tetO-OS<sup>AV</sup>KM #4 | 40, XY | 32 | 1 | 0 | 0 | 0 |
| tetO-OS<sup>AV</sup>KM #5 | 40, XY | 32 | 17 | 6 | 4 | 4 |
| tetO-OS<sup>AV</sup>KM #6 | 40, XY | 44 | 4 | 2 | 0 | 0 |
| epi-OSKM #1 | 40, XY | 33 | 15 | 11 | 6 | 5 |
| epi-OSKM #2 | 40, XY | 30 | 8 | 4 | 0 | 0 |
| epi-OSKM #3 | 40, XY | 42 | 0 | 0 | 0 | 0 |
| epi-OSKM #4 | 40, XY | 38 | 11 | 6 | 2 | 1 |
| epi-OS<sup>AV</sup>KM #1 | 40, XY | 55 | 10 | 2 | 0 | 0 |
| epi-OS<sup>AV</sup>KM #2 | 40, XY | 39 | 22 | 18 | 13 | 11 |
| epi-OS<sup>AV</sup>KM #3 | 40, XY | 29 | 17 | 17 | 10 | 10 |
| epi-OS<sup>AV</sup>KM #4 | 40, XY | 30 | 21 | 18 | 15 | 13 |
| 24h-tetO-OS<sup>*</sup>KM #1 | 40, XX | 12 | 5 | 3 | 2 | 0 |
| 24h-tetO-OS<sup>*</sup>KM #2 | 40, XY | 24 | 0 | 0 | 0 | 0 |
| tetO-OS<sup>*</sup>KM #1 | 40, XY | 24 | 12 | 9 | 8 | 0 |
| tetO-OS<sup>*</sup>KM #2 | 40, XY | 18 | 3 | 3 | 0 | 0 |
| epi-OKS #1 | 40, XY | 29 | 19 | 18 | 9 | 6 |
| epi-OKS #2 | 40, XY | 28 | 12 | 12 | 12 | 11 |
| epi-OKS #3 | 40, XY | 20 | 13 | 10 | 2 | 2 |
| epi-OKS #4 | 40, XY | 20 | 13 | 11 | 8 | 7 |
| epi-OKS<sup>*</sup> #1 | 40, XY | 30 | 23 | 17 | 7 | 5 |
| epi-OKS<sup>*</sup> #2 | 40, XY | 20 | 11 | 6 | 3 | 3 |
| epi-OKS<sup>*</sup> #3 | 40, XY | 30 | 12 | 10 | 6 | 5 |
| epi-OKS<sup>*</sup> #4 | 40, XY | 18 | 10 | 10 | 8 | 6 |
| epi-OKS<sup>*</sup> #5 | 40, XY | 19 | 8 | 8 | 6 | 6 |

### Engineered super-SOX enhances reprogramming across species

Next, we focused on Sox2-17, the most efficient chimeric Sox factor of this study, that includes 61V mutation along with three other structural elements of Sox17 (Fig. 1b-c). We cloned Sox2-17 into tet-inducible polycistronic OSKM or SKM reprogramming cassettes and confirmed comparable levels of expression using qPCR (Fig. S5a). Time-course experiments with restricted Dox-induction (Fig. 5a) showed that Sox2-17 dramatically enhances in the kinetics and efficiency of reprogramming (Fig. 5b). While at least three days of OSKM induction is needed to yield the first iPSC colonies, OS*KM carrying Sox2-17 gives rise to iPSC colonies after just 24h of induction—the fastest TF-based reprogramming reported to date. Clonally expanded 24h iPSCs lost methylation of *Nanog* and *Pou5f1* promoters, and acquired methylation of fibroblast-specific *Col1a1* promoter (Fig. S5b). 24h iPSCs could differentiate into all three germ layers in teratoma assays (Fig. S5c), contribute to chimeric mice (including the germline) (Fig. S5d), and even successfully generate live-born all-iPSC pups in 4N complementation assays (Fig. S5e, Table 1). When induced for just 3-4 days, OS*KM gave rise to 10 to 200 times as many colonies as OSKM, depending on the quality of starting fibroblasts (Fig. 5b-c). Sox2-17 could even generate mouse two-factor iPSCs with Klf4 alone (S*K cocktail) albeit with low efficiency, while Sox2, Sox2c17, or Sox17^EK^ could not (Fig. S5f). Clonal S*K iPSC lines displayed normal morphology (Fig. S5g) and PCR genotyping confirmed the identity of the two tested lines (Fig. S5h). S*K iPSCs stained positive for pluripotency markers Nanog and SSEA-1 (Fig. S5i), and gave rise to three germ layers in a teratoma assay (Fig. S5j). These data suggested that Sox2-17 requires shorter time, lower levels of overexpression, and a reduced number of additional factors to successfully induce pluripotency, which could be particularly beneficial for much less efficient integration-free reprogramming methods.

**Fig. 5.**
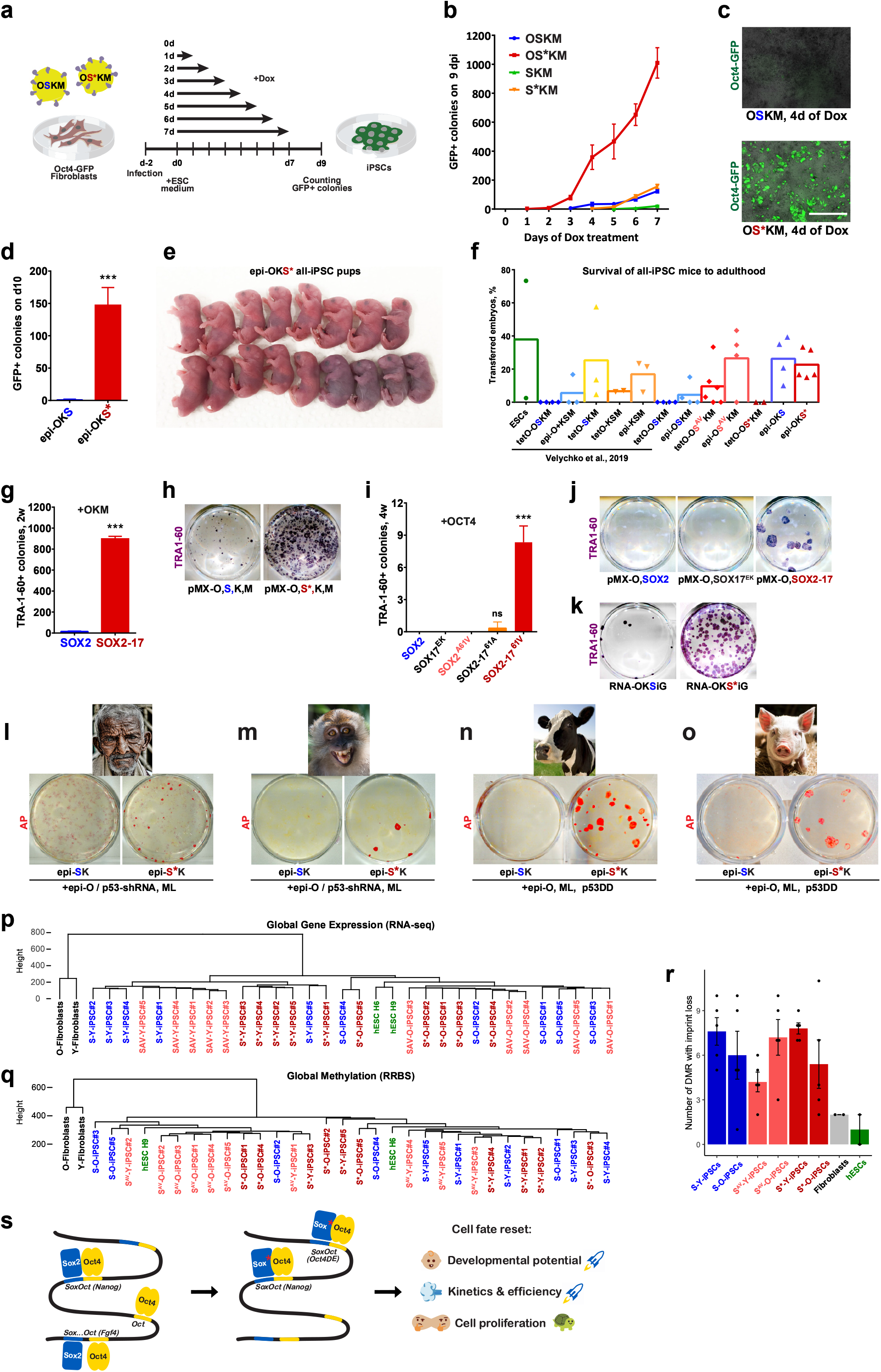
Sox2-17 enhances reprogramming in five species. **a,** Schematic representation of time course tet-inducible lentiviral reprogramming experiment. **b,** Time course reprogramming of Oct4-GFP MEFs induced with OSKM or SKM carrying either Sox2 (S) or Sox2-17 (S*) for the given number of days; the iPSC colonies were counted on 9 dpi. **c**, Representative brightfield and Oct4-GFP merged overview images of MEFs induced for 4 days with OSKM or OS*KM, images taken on 9 dpi, scale = 2 mm. **d**, Reprogramming of Oct4-GFP MEFs using episomal OKS (pCXLE-Oct4-P2A-KIf4-IRES-Sox) carrying either wild-type Sox2 or Sox2-17. 1.5×10^5^ cells were transfected with Fugene6. **e**, All-iPSC pups generated by the tetraploid (4N) complementation assay with epi-OKS* iPSC#l. 20 aggregates were transferred to two pseudopregnant CD-1 (white) females. **f**, Percentage of 4N-aggregated embryos that gave rise to adult healthy mice (survived till at least 3 months), including our previously published results^22^. Only XY lines were plotted. Data are represented as the mean of all tested lines. **g**, Reprogramming of human fetal fibroblasts (CRL-2097) with monocistronic retroviral OSKM carrying either wild-type SOX2 or SOX2^43–47,61,65–86^c17 (SOX2-17, S*). The volumes of viral supernatants were adjusted according to the qPCR titration. 10^4^ transduced cells were plated on a feeder layer of 6 well-plates. TRA1-60+ colonies were counted after two weeks of infection. Error bars represent SD; n = 3. **h**, Representative whole-well scan of (g). **i**, Two-factor (OS) reprogramming of human fibroblasts with monocistronic retroviral vectors. TRA1-60+ colonies were counted after four weeks of infection. Error bars represent SD; n = 3. **j**, Representative whole-well scan of **(i). k,** TRA1-60 staining of human fetal (CRL-2097) fibroblasts reprogrammed with self-replicating RNA (VEE) vectors carrying either OKSiG or OKS*iG reprogramming cassettes. **l-o**, Whole-well scans of alkaline phosphatase (AP) staining for episomal reprogramming of **(l)** 56-year-old human male dermal fibroblasts on day 25, (**m**) Cynomolgus macaque (*Macaca fascicularis*) fibroblasts on day 25, **(m)** bovine fetal fibroblasts on day 21, (**o**) porcine fetal fibroblasts on day 21 after nucleofection. **p-q**, Hierarchical clustering analysis of human ESC and integration-free iPSC lines derived from human newborn foreskin (young, Y) or 56-year-old human dermal (old, O) fibroblasts using episomal OSKML (pCXLE-OCT4+shP53, L-MYC-F2A-LIN28 and SOX-P2A-KLF4 carrying SOX2, SOX^AV^, or SOX2-17) based on: global gene expression (RNA-seq), TPM ≥1 (**p**) or global methylome (RRBS) (**q**). Clustering was based on Euclidean distance. **r**, Comparison of the number of genes that lost imprinting in 23 DMRs according to RRBS data. **s**, A model of the advantageous effects of highly cooperative Sox factors on cell fate reset.

We generated mouse episomal polycistronic Oct4-P2A-KIf4-IRES-Sox vectors carrying either wild-type Sox2 or Sox2-17 (epi-OKS and epi-OKS*, respectively) and confirmed the correct expression with western blot (Fig. S5k). Sox2-17 enhanced epi-OKS reprogramming of MEFs by a striking 150 times (Fig. 5d), giving rise to high-quality iPSCs that could generate all-iPSC mice with up to 77% efficiency (Fig. 5e, Table 1). Remarkably, all 10 tested integration-free epi-OKS and OKS* iPSC lines gave rise to healthy adult all-iPSC mice with the survival rate similarly high for both Sox2 and Sox2-17 (Fig. 5f, Table 1).

We tested a human version of SOX2-17 (Fig. 1c) for reprogramming human fibroblasts by delivery of retroviral monocistronic OSKM (Fig. 5g-h). The transduced cells were plated on inactivated feeders, and, after two weeks, stained for the human pluripotency marker TRA1-60. SOX2-17 gave rise to 56 times more iPSC colonies compared to SOX2: 8.9% and 0.16% overall reprogramming efficiency, respectively (Fig. 5g-h). We found that SOX2-17 can reprogram human cells even when combined with OCT4 alone (OS* cocktail), albeit with low efficiency (Fig. 5i-j). We showed that 61V within SOX2-17 is crucial for enabling two-factor reprogramming (Fig. 5i). In contrast, SOX17^EK^ did not give rise to a single human OS iPSC colony (Fig. 4i-j). Human clonal OS* iPSCs showed normal morphology (Fig. S5I), expressed pluripotency markers NANOG and TRA1-81 (Fig. S5m), and could differentiate into tissues of three germ layers in teratoma assays (Fig. S5n). Human self-replicating RNA^68^ OKS*iG vector encoding OCT4, KLF4, SOX2-17, and GLIS1 also generated about 50 times more TRA1-60+ colonies comparing to wild-type SOX2 (Fig. 5k). Moreover, using the RNA-OKS*iG vector, we have successfully generated and characterized integration-free iPSCs from dermal fibroblasts of a Parkinson’s disease patient, which in our hands could not be reprogrammed using the original vector (Schwamborn et al., manuscript in preparation).

We also cloned SOX2^AV^ and SOX2-17 into the episomal SOX2-F2A-KLF4 vector^69^, replacing the original F2A self-cleaving peptide with P2A to reduce formation of a fusion poly-protein^39^. Western blot confirmed equal expression and the correct cleavage of SOX2/SOX2^AV^/SOX2-17 and KLF4 (Fig. S5o). SOX2-17-P2A-KLF4 (S*K) combined with OCT4/shTP53, and L-MYC/LIN28 (OS*KML) vectors demonstrated much improved performance, particularly in reprogramming of aged human dermal fibroblasts, where reprogramming attempts using SOX2-carrying vector often did not yield any alkaline phosphatase-positive (AP^+^) colonies (Fig. 5I). To test if the super-SOX could facilitate reprogramming for other species, we tried our episomal vectors in generating iPSCs from cynomolgus macaque (*Macaca fascicularis*), an important non-human primate model^70–72^, for which ESCs have been derived^73,74^ but no transgene-independent iPSCs have been reported. The wild-type episomal OSKML vectors failed to give rise to a single AP^+^ colony during multiple attempts, while the same cocktail containing SOX2-17 gave rise to a few putative AP^+^ iPSC-like colonies (Fig. 5m). While most of hiPSC lines lose the episomal vectors before the third passage, only three out of 11 generated cynomolgus iPSC (ciPSC) lines completely lost the transgenes after three passages (Fig. S5p); two of the three lines had the correct chromosomal number (Fig. S5q). Integration-free ciPSCs displayed morphology similar to that of hiPSCs, except they were more prone to differentiation in the middle of colonies and could only be maintained on a feeder layer (Fig. S5r). They expressed pluripotency markers NANOG and OCT4 (Fig. S5s) and could differentiate into three germ layers in teratoma assays (Fig. S5t).

We also attempted to reprogram bovine and porcine fibroblasts using XAV939-containing bFGF-based media recently developed to derive and maintain cattle ESCs^75^. Episomal reprogramming using wild-type SOX2 failed in both species but the same cocktail containing SOX2-17 efficiently generated AP^+^ iPSC-like colonies for both cow (Fig. 5n) and pig (Fig. 5o). Bovine iPSCs (biPSCs) could even be generated without inhibiting p53 by the dominant negative mutant p53DD^76–78^, albeit with low efficiency (Fig. S5u). We established 12 of such epi-OS*KML biPSC lines (without p53DD), which all lost the episomal vectors by passage 6 (Fig. S5v). They could be passed at least 11 times, maintaining ESC-like morphology (Fig. S5w) and correct number of chromosomes (Fig. S5x), and stained positive for SOX2 and OCT4 (Fig. S5y).

To investigate the effects of SOX2^AV^ and SOX2-17 on the faithfulness of human reprogramming, we generated and characterized 30 hiPSC lines using episomal reprogramming of new-born foreskin (young, Y) and 56-year-old male dermal (old, O) fibroblasts. All the selected clonal iPSC lines tested were integration-free (Fig. S5z) with normal karyotypes (Fig. S5aa-ab). Hierarchical clustering of global gene expression based on RNA-seq showed that all iPSCs clustered far from fibroblasts and close to hESCs (Fig. 5p). The gene expression differences between the lines arose from the cell source more than the SOX factors used. We performed reduced representation bisulfite sequencing (RRBS)^79^ to analyze the methylome of hiPSCs. All lines clustered far from fibroblasts and close to hESCs (Fig. 5q). Loss of imprinting (LOI) is a common potentially cancerous^60,61^ irreversible^62^ epigenetic aberration afflicting iPSC technology for both mouse and human^63–66^. Multiple studies reported a correlation between LOI and poor developmental outcome of all-iPSC embryos in 4N complementation experiments^22,66,67,83,85–87^. We analyzed 23 differentially methylated regions (DMRs) represented in all samples and found that all lines including the original fibroblasts had different levels of LOI, with SOX2^AV^ hiPSCs derived from young fibroblasts and SOX2-17 hiPSCs derived from old fibroblasts exhibiting, on average, the lowest levels of LOI among the iPSC lines (Fig. 5r). We conclude that highly-cooperative Sox factors can allow high-efficiency high-quality reprogramming across species (Fig. 5s).

### Sox/Oct heterodimerization is at the core of naïve pluripotency

The ESC derivative of mouse pre-implantation inner cell mass and their iPSC counterparts grown in LIF containing media are called “naïve”^88^. Mouse naïve lines exhibit the highest developmental potential of all cultured cells: they readily contribute to chimeric animals, and some, but not all naïve lines are even capable of generating all-PSC mice. “Primed” PSCs of other species including human have much lower developmental potential and are more similar to mouse epiblast stem cells (mEpiSCs)^89^. Interestingly, *Oct4DE* is active in naïve but not primed PSCs^90–92^ and our highly-cooperative Sox factors dramatically increase the stability of Sox2/Oct4 heterodimer on *Oct4DE* element (Fig. 3f). Inspired by these results, we hypothesized that Sox/Oct cooperativity could be at the core of naïve pluripotency.

We analyzed a previously published time-course ATAC-seq dataset of naïve-to-primed transition samples generated for mouse ESCs (mESCs) upon exposure to bFGF media^93^. Spearman correlation of sequencing reads confirmed the reproducibility between replicates and showed that the most significant transition occurs between day one (d1) and day two (d2) of differentiation into epiblast-like cells (EpiLCs) (Figure 6a). Footprinting analysis using TOBIAS^57^ showed that the most depleted footprints on d1 were those of estrogen-related receptor (Esrr) and Klf factors (Figure 6b). This is consistent with previous studies showing that members of either family—Klf4 or Essrb—can convert mEpiSCs to naïve mESCs^94,95^. The d1 samples also had a significant depletion of Sox/Oct footprints (Figure 6b), which became the most depleted of all TF footprints between d1 and d2 of transition (Figure 6c).

**Fig. 6.**
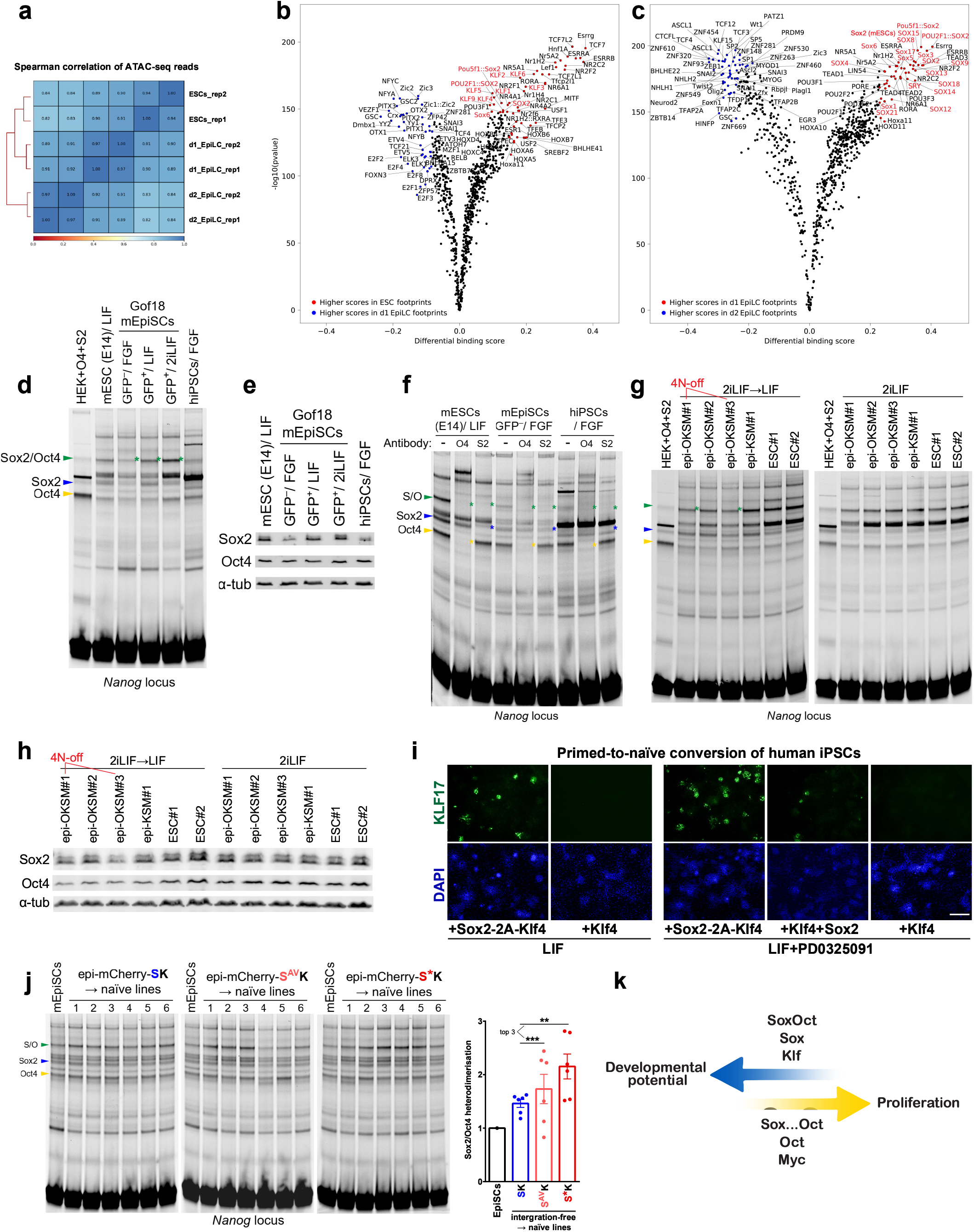
Sox/Oct cooperativity in high-versus low-grade pluripotency. **a**, Spearman correlation of time course ATAC-seq reads for naïve-to-primed differentiation mouse ESC samples^93^. **b-c**, TOBIAS footprinting analysis of (a) using MEME TF motif database of ESCs versus day one EpiLC samples (b), and day one versus day two EpiLC samples (c). **d**, EMSAs of whole-cell lysates showing endogenous levels of Sox2 and Oct4 that can bind *Nanog* promoter DNA element labeled with Cy5. Arrow heads indicate DNA bound by Oct4 (yellow), Sox2 (blue), or the heterodimer (green). All cells were grown without feeders. **e**, Western blot of lysates used in Fig. 6d. **f**, A supershift assay using anti-Oct4 and anti-Sox2 antibodies to confirm the identity of protein/DNA complexes in whole-cell lysate EMSAs. **g**, EMSAs of 6 mouse iPSC and ESC lines^22^, of which epi-OKSM1 and epi-OKSM3 were incapable of generating all-iPSC mice (4N-off). The lines were grown in 2iLIF on gelatin-coated plates, then split either in KSR-LIF media (left panel) or again in 2iLIF media (right panel). **h**, Western blot of lysates used in Fig. 6g. **i**, Immunostaining for naïve pluripotency marker KLF17 of primed-to-naïve converted human iPSCs on day 6 after transduction with constitutive lentiviral vectors (pHAGE2-EF1α). **j**, EMSAs of integration-free clonal mEpiSC lines primed-to-naïve converted using episomal mCherry-T2A-SOX-KLF4 vectors carrying wild-type SOX2, SOX2^AV^, or SOX2-17. All lines were converted and expanded on a feeder layer. The error bars represent SEM, statistical significance was calculated between top three out of six lines converted by each cocktail with Student’s t-test. **k**, SoxOct model of highgrade developmental reset.

We performed whole-cell lysate EMSAs on the *SoxOct* element of *Nanog* promoter to measure the levels of heterodimerization of naturally expressed Sox2 and Oct4 in different pluripotent lines: naïve mESCs grown in LIF-containing media, primed mEpiSCs carrying Gof18 reporter^96–98^ grown in bFGF-containing media, primed-to-naïve converted mEpiSCs grown in LIF or 2iLIF media (containing Mek and GSK-3 inhibitors, PD0325901 and CHIR99021^99^), and hiPSCs grown in conventional bFGF-based media (Fig. 6d). The EMSA results echoed our footprinting analysis: primed mEpiSCs had much lower levels of Sox2 and Oct4 capable of heterodimerizing compared to naïve mESCs, but mEpiSCs restored the heterodimerization level to the mESC level upon prime-to-naïve conversion (Fig. 6d). The heterodimerization (but also Oct4 alone binding) was further enhanced in the presence of 2i (Fig. 6d). The hiPSCs had low Sox2/Oct4 heterodimerization capacity, similar to mEpiSCs, but also higher levels of Oct4 alone binding. Western blot analysis showed that the lack of Sox2/Oct4 heterodimerization was due to lower Sox2 expression in both mouse and human primed cells; Oct4 expression levels were equal in mEpiSCs and mESC-LIF samples but higher in 2iLIF samples and in hiPSCs (Fig. 6e). We confirmed the identity of Sox2, Oct4, and Sox2/Oct4 bands in all tested lines using antibody super shift assays (Fig. 6f).

To find if Sox2/Oct4 heterodimerization could also explain the vast difference in developmental potential among iPSC lines, we tested six integration-free mouse PSC lines, which we characterized in detail before^22^. Two out of six iPSC lines (epi-OKSM#1 and epi-OKSM#3) were unable to generate full-term all-iPSC pups (4N-off, “poor quality”), while the other four (epi-OKSM#2, epi-KSM#1, mESC#l and mESC#2) could generate adult all-PSC mice^22^. The cells were first adopted to feeder-free conditions with several passages in 2iLIF media, then passed either with or without 2i and harvested for EMSA experiments. Of the cells grown in LIF-alone media, two developmentally incapable iPSC lines had the lowest capacity for Sox2/Oct4 heterodimerization (Fig. 6g) and the lowest Sox2 expression (Fig. 6h) among the six lines, demonstrating a pattern similar to mEpiSCs (Fig. 6d-e). However, the same lines passaged in 2iLIF showed no difference in either Sox2/Oct4 heterodimerization or Sox2 expression (Fig. 6g-h). This suggests that pluripotent cell lines could be stabilized at different levels of Sox2 expression: from Sox-high “very naïve” to different shades of “moderately naïve” and “primed” cultures with progressively lower developmental potency (Fig. 6a-h). We and others previously described capturing different grades of pluripotency ^97,100^, but the role of differential Sox2 expression and Sox2/Oct4 heterodimerization has not received much attention. The widely used 2i media seems to equalize the heterodimer content between different grades of naïve cells but fails to stably reprogram the cell fate: the mediocre Sox-low lines revert to being mediocre after 2i withdrawal (Fig. 6g-h). So, how can we stably “upgrade” the low-grade PSCs such as mEpiSCs or conventional pluripotent cultures in most other species including human?

mEpiSCs could be converted to naïve ESCs by overexpression of Klf4^95^. We attempted a similar conversion for human cells but found that Klf4 alone is not sufficient (Fig. 6i). Screening of different subsets of OSKM showed that overexpression of Sox2 and Klf4 (SK) is sufficient to convert conventional hiPSCs into dome-shaped colonies positive for human naïve pluripotency marker KLF17^101–104^ even in conventional KSR-LIF media in the absence of small molecule inhibitors (Fig. 6i). Similarly to SKM reprogramming in mice^22,25^, combining Sox2 and Klf4 in one bicistronic vector increased the number of KLF17^+^ colonies (Fig. 6i). Supplementing KSR-LIF media with Mek inhibitor enhanced the efficiency of the conversion (Fig. 6i). We used the SK cocktail for further experiments employing Gof18 mEpiSCs as a model. We wanted to achieve an integration-free pluripotency upgrade and to understand the role of Sox/Oct cooperativity in SK-based primed-to-naïve conversion. We cloned mCherry into the human episomal reprogramming vectors to generate mCherry-T2A-SOX-P2A-KLF4 carrying SOX2, SOX2^AV^, or SOX2-17. The episomal Cherry-SK vectors were delivered to mEpiSCs using lipofection, the cells were sorted for mCherry^+^ Gofl8^-^ on day 2 (Fig. S6a) and plated in KSR-LIF media on inactivated feeders. The majority of surviving cells formed dome-shaped colonies that were Gof18^+^/mCherry^-^ already on day 4 after plating (day 6 after transfection); six individual Gof18^+^/mCherry^-^ colonies were picked for each of the three cocktails and clonally expanded for further characterization (Fig. S6b). Regardless of the Sox version, the SK-converted lines exhibited big differences in Sox2/Oct4 heterodimerization determined by EMSAs (Fig. 6j), which correlated with differences in Sox2 expression determined by western blot (Fig. S6c), likely representing different grades of pluripotency. A transient overexpression of wild-type SK increased the Sox2/Oct4 heterodimerization by maximum of 62% compared to Gofl8^-^ mEpiSCs grown in the same conditions (Fig. 6j). Both highly-cooperative Sox factors appeared to be much more potent in resetting the Sox2/Oct4 heterodimerization: 3 out of 6 S^AV^K lines had 120-140% increase in heterodimerization, and two out of six S*K lines had a striking 180% boost (Fig. 6j). The integration-free S*K-converted naïve lines also had significantly lower propensity to spontaneously lose Gof18^+^ status during passaging (Fig. S6b).

These data suggest that the decrease of Sox2/Oct4 heterodimerization could be responsible for diminished developmental potential upon priming of PSCs in early development (Fig. 6k). SK reprogramming can reverse the process and highly-cooperative engineered Sox factors markedly promote this reset.

## DISCUSSION

iPSC technology remains inefficient particularly for non-murine cells and reprogramming often leads inconsistent iPSC quality^82–85^. A few alternative cocktails have been shown to improve the fidelity of reprogramming process in mice^22,66,67^ but failed to reprogram human cells, which appear to possess a stronger epigenetic barrier^105^. Perhaps the apparent barrier is merely a testimony to the inadequacy of the wild-type factor-based reprogramming machinery, which we currently employ. We and others have reported engineering enhanced reprogramming factors that increased the efficiency of mouse reprogramming^31,106–108^ but none of the studies showed a substantial gain of efficiency and faithfulness of iPSC generation beyond the mouse model. All alternative cocktails that improved the developmental potential of mouse iPSCs also decreased the efficiency of reprogramming, making them impractical even if they did work for humans^22,66,67^. We combined the structural elements of Sox2 and Sox17 to build a chimeric Sox2-17 that enhances reprogramming in five tested species: mouse, human, cynomolgus macaque, cow, and pig (Fig. 5, Fig. S5). Remarkably, just 24h of induction with our modified Yamanaka cocktail could generate mouse iPSCs that support full-term development, demonstrating the fastest TF-driven cell fate reset reported to date (Fig. 5a-b, Fig. S5a-e). We showed that Sox2-17 enhances mouse integration-free OKS reprogramming efficiency by 150 times generating consistently high-quality iPSCs (Fig. 5d-f, Table 1). A point mutation in Sox2, A61V, dramatically increased the developmental potential of mouse OSKM iPSCs, marked by both a higher rate of full-term development of all-iPSC embryos and the survival of the all-iPSC mice to maturity (Fig. 4g-i, Table 1). OSKM and OSK cocktails not only can induce a complete cell fate reset in the dish but also reverse aging of various animal tissues^4–9^. However, a very much expected lifespan extension for wild-type mice has not yet been demonstrated^8^. It is reasonable to assume, that engineered factors that induce iPSCs of higher quality (Fig. 4g-i, 5f) and support normal development of tissues to adulthood, even with likely background leaking of tet-inducible promoter^109^, could also outperform wild-type factors in reversing animal aging.

Notably, all the reprogramming cocktails that improved the developmental potential of mouse iPSCs, including OSKM compared to OKSM^85^, OSK (Fig. 5d-f)^67^ and SKM^22^ compared to OSKM, as well as OS^AV^KM compared to OSKM (Fig. 4g-i), also reduce the rate of cell division (Fig. 1i). Some cell-culture interventions that improve the quality of iPSCs also inhibit cell division^110^. Therefore, it is tempting to theorize that the secret of a perfect reset lies in limiting cell proliferation (Fig. 6k). This can be achieved by I) enhancing Oct/Sox cooperativity, as in OS^AV^KM reprogramming; II) increasing the Sox2 to Oct4 ratio, as in SKM reprogramming, but also in Oct4 heterozygous knockout ESCs^111^; III) omitting or reducing the level Myc, as in OKS (Fig. 5f, Fig. S5k)^67^ and in OSKM compared to OKSM cassette^85^, but also in Myc-depleted ESCs^112^; IV) reducing the overexpression levels of reprogramming factors, as in episomal versus viral vectors (Fig. 5f)^22^. Admittedly, following the episomal reprogramming protocols, we used the construct carrying shRNA against TP53^69^ to generate human epi-OSKML iPSC lines for this study, which knocks down the major tumor suppressor and boosts cell proliferation. This could have caused loss of genetic imprints in our human iPSC lines (Fig. 5r). Considering the mouse developmental potential results presented here, knocking down our genome guardian, is likely detrimental to the quality of iPSCs and should be avoided, just as with Myc (Fig. 5f).

Many studies addressed uniqueness of Oct4 among POU factors by domain swapping and mutagenesis^20,39,48,106,113–116^. We and others determined the importance of the Oct4-POU_S_, POU_HD_ linker, and CTD domains. Here, we report that A61V swap at the Sox/Oct interface of Sox2, increases the stability of the Sox/Oct complex on canonical *SoxOct* motifs that control pluripotency genes. A61V mutation, especially when combined with the stronger Sox17 C-terminal transactivator, can rescue the otherwise disabling deletions of the Oct4 linker, POU_HD_ NTD, or CTD domains. Oct4-POU_S_ domain that contains the Sox/Oct interface (Fig. 2) proved to be the most crucial for reprogramming (Fig. 3a-c). The 17 residue-long linker peptide connects the two DNA-binding subdomains of Oct4-POU domain. All the resolved POU structures so far suggest that the linker is not directly involved in the DNA-binding (Fig. 2a-b), yet it appears to be important for both reprogramming to pluripotency and for normal development^41,44,117^. Until now, the effects of linker mutations on Sox/Oct stability have not been tested. Here we found that Oct4 linker mutants, such as L80A or a complete replacement with flexible poly-glycine peptides, reduce stability of the heterodimers on DNA. Sox2^AV^ can rescue the ability of the linker mutants to heterodimerize and to induce pluripotency (Fig. 1f, 3a, S3k). Similarly, Sox2^AV^ enables heterodimerization and reprogramming with tissue-specific POU factors (Fig. 1g-h and S3b, k). Our molecular dynamics simulation (MDS) revealed that A61V enhances the hydrophobic interaction of HMG and POU_S_ domains of Sox/Oct heterodimer on canonical *SoxOct* motif, which in the case of Oct4, likely exist in multiple configurations (Fig. 2, S2). The negatively charged E78 and E82 of Oct4-linker have the propensity to form salt bridges with positively charged 57K and 50R of Sox2 (Fig. 2c, e, Fig. S2 a-b). Concordantly, Sox2^K57E^ mutant disrupts Sox2/Oct4 heterodimerization on canonical *SoxOct* motifs but not on *Fgf4*, which disables induction of pluripotency^36,46^. Our experimental data showed that Sox2^AV^ and Sox2-17 enhanced cooperativity with wild-type Oct4 on the *Nanog* promoter, *Oct4DE, HoxB1*, and *Utf1* enhancer loci, all containing canonical *SoxOct* motifs (Fig. 3, Fig. S3). Sox2^AV^ also enhanced heterodimerization on a nucleosomal *SoxOct* element (Fig. 4a) and in early reprogramming samples *in situ* (Fig. 4b-f). However, consistent with the model prediction (Fig. 2f), neither Sox2^AV^ nor Sox2-17 changed cooperativity on the *Fgf4* enhancer motif (Fig. 3f, Fig. S3j, Fig. 4e), which has three additional base pairs spaced between Sox and Oct elements^46^. Fgf4 regulates cell proliferation and differentiation in the epiblast^118,119^, and inhibiting the Fgf4 pathway with PD0325901 facilitates maintenance of naïve PSCs^119^.

Mouse naïve PSCs are the most developmentally potent cultures we currently have. Mice likely evolved (or preserved) the unusual stability of their naïve pluripotency fate to enable a blastocyst-stage embryonic arrest, known as diapause^120^. Extraordinary developmental potential of mouse naïve PSCs as well as increased capacity of naïve cells for homologous recombination repair, has allowed for unprecedented genetic engineering of this species^121^. Mouse remains the only species for which generation of all-iPSCs animals has been reported^122–124^; and germline competence has been reported only for mouse and male rat^125^, highlighting the limitations of today’s technologies. Naïve PSCs have lower Myc levels compared to primed cells^97,99^, which could be a direct effect of high levels of Sox2^126^ in naïve cells (Fig. 6e). Here, we demonstrated that Sox2/Oct4 heterodimerization is reduced during naïve-to-primed differentiation (Fig. 6b-c) and is restored during primed-to-naïve conversion (Fig. 6d). Moreover, the loss of Sox2/Oct4 heterodimerization also explains the diminished developmental potential in at least some 4N-off OSKM miPSCs grown in naïve media (Fig. 6d-h). The Sox/Oct model of developmental reset presented here (Fig. 6k) echoes other studies placing Sox2 at the top of pluripotency network hierarchy^21,28,29,127–131^. In early animal development, unidirectionality is likely achieved by a negative feedback loop limiting the return to a high-Sox2 state^132^. SK reprogramming can reset the high level of Sox2 expression (Fig. S6c), which restores Sox2/Oct4 heterodimerization (Fig. 6j) and activates naïve pluripotency markers (Fig. 6i, S6b). The specific enhancement of S and SL, but not DS configuration by A61V (Fig. 2,3f) could be the key for the enhanced developmental potential of OS^AV^KM iPSCs (Fig. 4g-i) and for the superior ability of highly cooperative Sox TFs to restore the Sox2/Oct4 heterodimerization in primed cells (Fig. 6j). Additional targeted disruption of DS binding on the *Fgf4* enhancer could potentially further improve the super-SOX-based developmental reset technology.

In animal evolutionary tree suggests that of the Sox2/Oct4 couple, in the beginning there was Sox2. Both 50R and 57K residues are already present in sponges, where SoxB TFs control early embryogenesis^133^; V at position 61 could also be found in Sox factors of these most primitive animals. 57K and 61A of Sox2 are conserved in hydrozoans, where SoxB genes are expressed in stem cells that give rise to neuroectoderm (default) and germline^134–136^. POU5 factors emerged much later in the evolutionary tree - it is an innovation of vertebrates^137^, where POU5 TF (or TFs) cooperate with Sox2 to control early development^138,139^. Interestingly, while the linker is the least conserved part of POU domains, the negative charges at positions 78 and 82 of POU5-linker are already present in jawless hagfish^137^. Our work demonstrates that the most significant feature that distinguishes Oct4 from other POU factors is its ability to form a stable heterodimer with Sox2, which had already been in control of early embryogenesis in lower animals.

Interestingly, it is well-known that mouse female pluripotent stem cells have lower developmental potential compared to their male counterparts^140,141^. Our model suggests that the reason for the increased developmental potential in male versus female pluripotent lines could be the expression of Sex determining region of Y (Sry) in XY lines^142^. The Sry binding motif is similar to that of Sox2 and, as in case of other Sox TFs, Sry footprints were depleted upon priming of naïve cells (Fig. 6c). The high-SOX hypothesis could also explain the increased survival of male versus female embryos in mammals^143,144^ and the higher occurrence of glioblastoma and other types of Sox-driven cancers in men^145^. The prospect that the mechanisms uncovered here could allow us to derive developmentally-capable PSCs across species is intriguing, yet, our new understanding of developmental reset will have ramifications beyond the pluripotency field.

A small number of cells of the inner cell mass can make a disproportionally high contribution to animal development^146^. Therefore, it is likely that the developmental potential of a given pluripotent line is determined by a Sox-high subpopulation and not the average Sox2 expression determined in our bulk EMSA and western blot experiments. More work is needed to address the heterogeneity within pluripotent lines, the mechanisms regulating Sox2 expression and functionality, and the possibilities to enrich and stabilize the Sox-high pluripotent cell cultures with presumably upgraded developmental potential. We cannot exclude that highly cooperative or an excess of Sox factors participate in the developmental reset in other ways, besides enhancing Sox/Oct heterodimerization, e.g. cellular remodeling by recruiting the antagonist of aging Parp1^147^ or silencing retroviral elements^22^.

Our engineered super-SOX harvests the beneficial naturally evolved structural elements of two major development regulators, Sox2 and Sox17. It is likely that even more efficient reprogramming factors could be built by means of rational engineering and directed evolution^106,107^. Our data suggest that enhancing cooperativity between key co-factors should be one of the goals of future designers.

## METHODS

### Mice

All mice used were bred and housed at the mouse facility of the Max Planck Institute in Münster. Animal handling was in accordance with MPI animal protection guidelines.

### Vector construction

The pMX-Sox2/Sox17 chimeric TF vectors were based on Addgene ID 13367^2^ and the tet-inducible pHAGE2-tetO-Oct4-P2A-Sox2-17-T2A-Klf4-E2A-cMyc (OS*KM) and pHAGE-tetO-Sox2-17-T2A-Klf4-E2A-cMyc (S*KM) vectors were based on Addgene ID 136551 and 136541, respectively^22^.

The self-replicating RNA vector T7-VEE-OKS*iG was based on Addgene ID 58974^68^. The mouse episomal vectors pCXLE-Oct4-P2A-KIf4-IRES-Sox2 (OKS) and pCXLE-Oct4-P2A-KIf4-IRES-Sox2-17 (OKS*), and human episomal vectors pCXLE-SOX2-P2A-KLF4 (SK), pCXLE-SOX2^AV^-P2A-KLF4 (S^AV^K), pCXLE-SOX2-17-P2A-KLF4 (S*K), pCXLE-mCherry-E2A-SOX2-P2A-KLF4 were based on Addgene ID 27078^78^, except the inefficient self-cleaving peptide F2A was replaced with P2A to avoid protein fusion. pCXLE plasmids showed much better yields when grown in Stbl2 competent E. coli (Invitrogen).

The mouse and human protein sequences of Sox2^AV^ and Sox2-17 were (HMG-box domains are uppercase, Sox17 parts are in bold):

>Mouse Sox2^A61V^ mynmmetelkppgpqqasgggggggnataaatggnqknspDRVKRPMNAFMVWSRGQRRKMAQENPKM HNSEISKRLGAEWKLLSETEKRPFIDEAKRLR**V**LHMKEHPDYKYRPRRKTKTLMKKDKytlpgglla pggnsmasgvgvgaglgagvnqrmdsyahmngwsngsysmmqeqlgypqhpglnahgaaqmqpmhrydvsalqynsm tssqtymngsptysmsysqqgtpgmalgsmgswkseasssppwtssshsrapcqagdlrdmismylpgaevpepaapsrlh maqhyqsgpvpgtaingtlplshm
>Human SOX2^A61V^ mynmmetelkppgpqqtsgggggnstaaaaggnqknspDRVKRPMNAFMVWSRGQRRKMAQENPKMHN SEISKRLGAEWKLLSETEKRPFIDEAKRLR**V**LHMKEHPDYKYRPRRKTKTLMKKDKytlpggllapgg nsmasgvgvgaglgagvnqrmdsyahmngwsngsysmmqdqlgypqhpglnahgaaqmqpmhrydvsalqynsmtss qtymngsptysmsysqqgtpgmalgsmgswkseasssppwtssshsrapcqagdlrdmismylpgaevpepaapsrlhms qhyqsgpvpgtaingtlplshm
>Mouse Sox2-17 mynmmetelkppgpqqasgggggggnataaatggnqknspDRVKRPMNAFMVWSRGQRRKMAQENPKM HNSEISKRLGAEWK**A**L**TLA**EKRPFIDEAKRLR**V**LHM**QD**HP**N**YKYRPRR**RKQVKRMKRVeggflh alvepqagalgpeggrvamdglglpfpepgypagpplmsphmgphyrdcqglgapaldgyplptpdtspldgveqdp affaaplpgdcpaagtytyapvsdyavsveppagpmrvgpdpsgpampgilappsalhlyygamgspaasagrgfh aqpqqplqpqapppppqqqhpahgpgqpspppealpcrdgtesnqptellgevdrtefeqylpfvykpemglpyqgh deg vn Isdshgaisswsdassavyycny pd i**
>Human SOX2-17 mynmmetelkppgpqqtsgggggnstaaaaggnqknspDRVKRPMNAFMVWSRGQRRKMAQENPKMHN SEISKRLGAEWK**A**L**TLA**EKRPFIDEAKRLR**V**LHM**QD**HP**N**YKYRPRR**RKQVKRLKRVeggflhgla epqaaalgpeggrvamdglglqfpeqgfpagppllpphmgghyrdcqslgappldgyplptpdtspldgvdpdpaffa apmpgdcpaagtysyaqvsdyagppeppagpmhprlgpepagpsipgllappsalhvyygamgspgagggrgfq mqpqhqhqhqhqhhppgpgqpspppealpcrdgtdpsqpaellgevdrtefeqylhfvckpemglpyqghdsgvnl pdshgaissvvsdassavyycnypdv**

All the relevant plasmids will be available from Addgene.

### Cell culture

HEK293 cells were cultured in low-glucose DMEM (Sigma) supplemented with 10% FBS (Capricorn Scientific), 1% Glutamax, 1% penicillin-streptomycin, 1% nonessential amino acids (all from Sigma). Mouse, human, cynomolgus and porcine fibroblasts were cultured in high-glucose DMEM (Sigma) supplemented with 15% FBS, 1% Glutamax, 1% penicillin-streptomycin, 1% nonessential amino acids (NEAA), 1% sodium pyruvate (Sigma), 1% β-mercaptoethanol (Gibco); bovine fibroblasts were cultured in 50:50 DMEM/F12 (Gibco) and IMDM (with HEPES, Cytiva) with 15% FBS and the same supplements; 5 ng/ml of human bFGF (Peprotech) was used to improve cynomolgus, bovine, and porcine fibroblast cultures.

Mouse naïve pluripotent stem cells were grown in KSR-based mouse embryonic stem cell (mESC) media: high-glucose DMEM medium supplemented with 15% KSR (Invitrogen), 1% Glutamax, 1% NEAA, 1% penicillin-streptomycin, 1% β-mercaptoethanol, and 20 ng/ml human recombinant LIF (purified in-house) on Mitomycin C-inactivated C3H MEF feeder layer. For 4N-complementation experiments the KSR-LIF media was supplemented with 2i (1 mM PD0325901 and 3 mM CHIR99021) for one passage. Mouse Gof18 GFP-E3 epiblast stem cells (EpiSCs)^148^ were cultured in Stem Flex media (Gibco) on FBS-coated dishes.

Human PSCs were cultured in hESC media: either in DMEM/F12 supplemented with 15% KSR, 1% Glutamax, 1% NEAA, 1% penicillin-streptomycin, 1% β-mercaptoethanol and 5 ng/ml bFGF or in StemFlex media (Gibco) on Matrigel-coated dishes (Corning) or on Mitomycin C-inactivated CF1 MEF feeder layer. Cynomolgus iPSCs were cultured in StemFlex media on Mitomycin C-inactivated CF1 MEF feeder layer. Bovine and porcine iPSCs were derived and cultured in StemFlex media supplemented with 2 μM XAV939 (Sigma) on Mitomycin C-inactivated CF1 MEF feeder layer in a hypoxic 5% O_2_, 5% CO_2_ incubator at 37°C; other cells were cultured in normoxic conditions.

Pluripotent stem cells of all five species were passed using Accutase (Sigma). 10 μM Rho-associated kinase inhibitor (Y-27632, Abcam) was added for the first 24h after passaging of primed PSCs of all five species (extended to 48h for mouse EpiSCs). The cells were routinely tested for Mycoplasma contamination and tested negative.

### iPSC generation

Mouse reprogramming experiments were done as described before^22,39^. Briefly, for retrovirus production monocistronic pMX-Oct4, Sox, and Klf4 vectors were co-transfected with pCL-Eco (Addgene ID 12371)^149^ in HEK293 cells with FuGENE6 (Promega) using low volume transfection protocol (Steffen et al., 2017). For lentivirus production pHAGE2-tetO vectors were co-transfected with PAX2 and VSV. The viral supernatants were harvested after two and three days, filtered (Millex-HV 0.45 μm; Millipore) aliquoted and stored at −80°C. For reprogramming, Oct4-GFP MEFs (OG2 or Rosa26rtTA-Gof18) were plated on gelatin-coated 12-well plates at 3×10^4^ cells per well in fibroblast media. A few hours later the cells were infected with titer-adjusted volumes of each viral supernatant supplemented with 6 μg/ml (final concentration) of protamine sulfate (Sigma). After two days, the media was replaced with mouse ESC media. For mouse tet-inducible reprogramming, the cells were treated with Dox for 10 days (same as^22^), unless otherwise stated. Because all the reprogramming experiments were treated equaly, enhanced kinetics of OS*KM reprogramming resulted in mature tetO-OS*KM expressing the Myc-containing transgene for much longer compared to tetO-OSKM or tetO-OS^AV^KM iPSCs that emerged later in the 10-day course. This explains the poor quality of tetO-OS*KM versus tetO-OS^AV^KM iPSCs.

For human retroviral reprogramming, 48h after infection, the transduced cells were split on a CF1 feeder layer at 10^4^ per 6-well plate. After one week, fibroblast media was changed to hESC media. For mouse episomal reprogramming 10^5^ of Oct4-GFP (Rosa26TA-Gof18) MEFs were plated on gelatin-covered 6-well plates overnight, and transfected with 1.5 μg of pCXLE-OKS or OKS* combined with 0.5 μg of pCXWB-EBNA1 (Addgene ID 37624) with FuGENE6.

Human self-replicating RNA-based reprogramming was performed as previously described^68^. Briefly, the T7-VEE constructs were digested with Mlul and then *in vitro* transcribed using RiboMAX Large Scale RNA Production System Kit (Promega). The transcripts were 2’-O-methylated, capped, and poly(A)-tailed using respective CELLSCRIPT kits following the manufacturer’s protocol. For reprogramming, 1 μg of RNA replicons were transfected into 10^5^ fibroblasts on 6-well plates using RiboJuice (Sigma) in the presence of 100 ng/ml B18R (Promega). The media was supplemented with 0.5 mM VPA, 5 μM EPZ to enhance the very inefficient RNA-based reprogramming. The reprogramming worked more efficiently when no puromycin selection was used. After two weeks, the cells were sorted for TRA-1-60 and plated on a CF1 feeder layer in human ESC media without B18R.

Human and cynomolgus episomal reprogramming was done as previously described^150^. Briefly, 5×10^5^ human newborn foreskin fibroblasts (Young, Y)^151^, 56-year-old male dermal fibroblasts (Old, O, AG04148), or cynomolgus (MHH Hannover) were nucleofected with 3 μg of plasmid DNA mix: pCXLE-SOX2-P2A-KLF4 or pCXLE-SOX2-17-P2A (made for this study), pCXLE-L-MYC-F2A-LIN28 (ML, Addgene ID 27080), pCXLE-hOCT4-shTP53 (Addgene ID 27077), pCXWB-EBNA1 (Addgene ID 37624) using Lonza NHDF Nucleofector kit (U-23 program), and plated in ROCKi-containing fibroblast media on a CF1 feeder layer at different densities.

For cattle reprogramming, 10^6^ bovine fetal fibroblasts (GOF 451-1)^152^ or porcine fetal fibroblasts ^153^ were nucleofected with 6 μg of plasmid DNA mix: pCXLE-SOX2-P2A-KLF4 or pCXLE-SOX2-17-P2A, pCXLE-L-MYC-F2A-LIN28, pCXLE-hOCT4 (Addgene ID 27076), pCXLE-p53DD (Addgene ID 41859), and pCXWB-EBNA1 using the human protocol. For bovine reprogramming, pCXLE-p53DD could be omitted.

The virus supernatant volumes were adjusted according to qPCR titration using common WPRE or 3’UTR primers normalized to Rpl37a^22^. All the tetO lines were screened for promoter leaking, only those with minimal leaking were selected for characterization. The newly generated iPSC lines (mouse, human, cynomolgus, and cow) were karyotyped using DAPI staining of metaphase spreads, only the lines with correct chromosomal numbers were selected for characterization. As we reported before ^22^, no difference in aneuploidy occurrence was observed between different cocktails. Similar to other studies, we only tested the quality of male iPSCs for this work.

### Primed-to-naïve conversion

For primed-to-naïve conversion (pluripotency upgrade), human iPSCs were transduced with monocistronic or polycistronic pHAGE2-EF1α lentiviral vectors carrying reprogramming factors. After two days, the cells were passed at low density (10^3^ cells per 24-well plate) on an inactivated C3H feeder layer in mESC media supplemented with ROCKi with or without 2i. 24h later the media was changed to mESC media without ROCKi with or without 2i. Six days after passing, the cells were fixed and stained for KLF17 (HPA024629, ATLAS, 1:500). PD0325901, but not CHIR99021 or PD0325901+CHIR99021 (2i) increased the number of KLF17^+^ colonies. For integration-free pluripotency upgrade, 3×10^5^ of GFP-negative Gof18 E3 mouse epiblast stem cells (mEpiSC)^148^ cells were seeded in StemFlex+ROCKi media on FBS-coated 12-well plates and simultaneously transfected with 2 μg of episomal pCXLE-mCherry, pCXLE-mCherry-T2A-SOX2-P2A-KLF4, pCXLE-mCherry-T2A-SOX2^AV^-P2A-KLF4, or pCXLE-mCherry-T2A-SOX2-17-P2A-KLF4 using 4μL of Lipofectamine Stem Reagent (Invitrogen) according to manufacturer’s instructions; after 48h the cells were sorted for mCherry and plated in mouse ESC media +ROCKi on an inactivated C3H feeder layer at 10^4^ per 12-well plate. ~30% of sorted cells survived; of those ~50% of SK/S^AV^K/S*K-transfected colonies grew dome-shaped and were GFP+/mCherry^-^ already on day 4 after passing. The GFP^+^ colonies were picked and clonally expanded for further characterization.

### Tetraploid (4N) complementation assay

1. Preparation of tetraploid embryos Super ovulated B6C3F1 females were mated with CD1 males. E1.5 embryos at the two-cell stage are flushed from the oviducts and collected in M2 medium. After the equilibration in fusion solution (0.3 M D-mannitol, 50 μM CaCl2, 0.3% BSA (Sigma)) 50-75 embryos are placed between the electrodes of a 250 μm gap electrode chamber (BLS Ltd.) containing 0.3 M mannitol with 0.3% BSA and fused with a Cellfusion CF-150/B apparatus (BLS Ltd.) with 0.5 mm Microslide (BTX-450). An initial electrical field of 2V is applied to the embryos followed by one peak pulses of 60V for 50 μs. Embryos are transferred back into KSOM-aa medium and immediately into a 37°C incubator with 5% CO_2_. Embryos are observed for fusion after 15 to 60 minutes. The fused tetraploid embryos are cultured for 24h to the 4-cell stage under the same conditions.
2. Aggregation of iPSCs with zona-free embryos
  1. Preparation of aggregation plates for mouse embryos chimera production 1h before aggregation: Using a KSOM medium filled 100μl-pipette, make 4 rows of microdrops (roughly 3mm in diameter) in a 35mm dish (Falcon, Cat. No. 35-3001), two drops in the first and fourth, five drops in the second and third rows. Cover the whole plate with paraffin oil. Sterilize the aggregation needle (BLS Ltd.) with 70% ethanol. Press the aggregation needle into the plastic through the paraffin oil and culture medium, while making a circle movement to create a tiny scoop of about 300 μm in diameter with a clear smooth wall. Six to ten holes can be made within each droplet.
  2. iPSCs are aggregated and cultured with denuded 4-cell stage mouse tetraploid embryos as reported with a slight modification^154^: Clumps of loosely connected iPSCs (15-20 cells in each) from short trypsin-treated day two iPSC cultures were chosen and transferred into microdrops of KSOM medium under mineral oil; each clump is placed in a depression in the microdrop. Meanwhile, batches of 30-50 embryos were briefly incubated in acidified Tyrode’s solution ^155^ until dissolution of their zona pellucida. Two embryos were place on the iPSC clump. All aggregates are assembled in this manner, and cultured overnight at 37°C, 5% CO_2_. After 24h of culture, the majority of aggregates have formed blastocysts. Ten to fourteen embryos were transferred into one uterine horn of each 2.5 days post coitum, pseudopregnant CD1 female that had been mated with vasectomized males. For Cesarean Section, recipient mothers were sacrificed at E 19.5 and pups were quickly removed. Newborns that were alive and respirating were cross fostered to lactating females.

### Mammalian cell overexpression and whole-cell lysate (WCL) generation

HEK293 cells cultured on 10cm dishes were transfected with 10 μg of pLVTHM or pHAGE2 vectors under the control of an EF1α promoter and containing the wild-type or mutant versions of Oct4 or Sox2 with Fugene6 (Promega) using a low volume protocol (Steffen et al., 2017). Three days after transfection, the cells were dissociated from the plate using Accutase (Sigma), collected, counted, and washed with PBS. WCLs were generated by five cycles of freeze-thawing pellets resuspended in 12.5 μL per million cells in lysis buffer (20 mM HEPES-KOH pH 7.8, 150 mM NaCl, 0.2 mM EDTA pH 8, 25% glycerol, 1 mM DTT, and cOmplete™ protease inhibitor cocktail (Merck). After disruption, lysates were spun at 14k RCF at 4°C for 10 min. After centrifugation, pellets were discarded and the supernatants transferred to a new tube for further analysis. Protein concentrations were estimated by diluting samples in 0.1% SDS solution, measuring A_230_ and A_260_, and applying the equation:

Cone. (μg/μL) = (0.183*A_230_-0.075*A_260_)*dilution factor

All samples were diluted to 1 μg/μL, aliquoted, snap frozen, and stored at −80°C. Western blots were run to compare expression levels between mutants. Expression was evaluated by Quantity One® (v4.6.7, Bio-Rad) densitometry to adjust for equal amounts of expression using WCL of untransfected cells to maintain total protein content, when necessary.

### Western blot analysis

5-10 μg of total protein was combined with Laemmli sample buffer, heated, and loaded onto 12% mini SDS-polyacrylamide gel (SDS-PAG) using the Towbin buffer system^156^. Gels were run initially at 15V for 15 minutes to load samples into the stacking gel and then 50 V for 30-60 minutes to resolve the proteins of interest. Samples were transferred to Immobilin®-FL PVDF membranes (Merck Millipore Ltd.) at 4°C under 300V for 2h. Membranes were blocked for one hour at room temperature in 5% skim milk (Sigma) dissolved in PBS with 0.1% Tween-20 (PBS-T) and incubated overnight at 4°C with rotation in the primary antibody diluted in blocking solution. The following day the membrane was washed three times in PBS-T and then incubated in secondary antibody diluted in blocking solution for one hour at 25°C. The following antibodies were used: polyclonal goat anti-Oct4 N-19 (sc-8628, Santa Cruz Biotechnology) or monoclonal mouse anti-Oct4 (611203, BD Biosciences), polyclonal goat anti-Sox2 (sc-17320 from Santa Cruz Biotechnology or GT51098 from Neuromics), monoclonal mouse anti-alpha tubulin (T6199, Sigma), 647-conjugated anti-goat (Alexafluor), and 647-conjugated anti-mouse (Alexafluor). Western blots signal was detected using Fujifilm FLA-9000 fluorescence scanner (Fujifilm).

### Insect cell expression and protein purification

The coding sequence of full-length *Mus musculus* Sox2 or Sox2^AV^ was cloned into pCoofy27 plasmid with an N-terminal 6xHis-tag using SLIC as previously described: forward primer 3C, reverse primer ccdB^157^. Plasmids were then transformed into DHIOEMBacY (a gift from Dr. Imre Berger) for baculovirus plasmid DNA amplification^158^. Bacmids were purified using Macherey-Nagel Xtra BAC100 (Düren) and then transfected into a suspension of Sf9 cells at 0.8×10^6^ cells/mL grown in serum-free EX-CELL® 420 medium containing L-glutamine (Sigma) and incubated at 26°C with shaking for virus production. Cells were monitored daily for increased cell size and GFP fluorescence. Once ~90% of cells were GFP+, viral suspensions were spun down and then filtered through 0.22 mm. Viral supernatants were expanded once before used for infection, filtered aliquots were stored at −80°C.

Optimal protein expression conditions were determined empirically. Mid-log phase High Five™ insect cells were split to 10^6^ cells/mL in 2 L and then infected with 10-12 mL of P1 baculovirus from previous steps per liter of cells. Following incubation at 28°C for 96h with shaking, cell pellets were collected by centrifugation. Pellets were resuspended in lysis buffer (20 mM HEPES pH 7.5, 300 mM NaCl, 30 mM lmidazole, 5% glycerol, 0.1% Triton X-100, cOmplete™ protease inhibitor cocktail (Merck), and 1 mM DTT), frozen and thawed once, then sonicated at 4°C using a probe sonicator (Bandelin Sonopuls, Bandelin Eletronics). Pellets were resuspended in inclusion body wash buffer (20 mM HEPES pH 7.5, 200 mM NaCl, 1 mM EDTA, 1% Triton X-100, cOmplete™ protease inhibitor cocktail (Merck), and 1 mM DTT) and subject to four cycles of Dounce homogenization followed by centrifugation for 20 min. at 18k RCF and 4°C, twice with inclusion body wash buffer and twice in buffer without Triton X-100. The final pellet was cut twice in DMSO and then incubated for 30 min at 25°C. Unfolding buffer (7 M guanidine hydrochloride, 20 mM Tris-HCl pH 7.5, 5 mM DTT) was added to the pellet and incubated while rotating for 1h at 25°C. Nickel Sepharose slurry (GE Healthcare) was washed and equilibrated in binding buffer, then supernatant was added and incubated at 4°C overnight with rotation. Proteins were eluted using the unfolding buffer with additional 500 mM imidazole. Eluate fractions were checked with SDS-PAGE and relevant fractions were pooled. Using 7 kDa molecular weight cut off (MWCO) dialysis tubing, pooled fractions were dialyzed for three buffer changes of at least six h for each volume of refolding buffer at 4°C (7 M urea, 20 mM Na Acetate pH 5.2, 200 mM NaCl, 1 mM EDTA, and 5 mM DTT). Following centrifugation to remove any insoluble material, the supernatant was dialyzed (7 kDa MWCO) in refolding buffer with decreasing amounts of urea: 1 hour 6 M urea, 2h 4 M, 2h 2 M, and 1 hour in size exclusion chromatography (SEC) buffer (50 mM Tris-HCl pH 7.4, 150 mM NaCl, 1 mM EDTA, 5% glycerol). Eluate was centrifuged to remove any precipitate before loading onto HiLoad 16/60 Superdex 200 SEC column (GE Healthcare).

The coding sequence for full-length Oct4 from *M. musculus* was cloned into the pOPIN expression vector using the SLIC method and Phusion Flash High-Fidelity PCR Master Mix (Finnzymes/New England Biolabs). SLIC reactions were then transformed into One Shot™ OmniMAC™ 2 T1® Chemically Competent *E. coli* (ThermoFisher Scientific). After sequencing, the pOPIN-cHis-Oct4 construct was co-transfected with flashBACULTRA™ bacmid DNA (Oxford Expression Technologies) into Sf9 cells (ThermoFisher Scientific) using Cellfectin II® (ThermoFisher Scientific) to generate recombinant baculovirus. Mid-log phase Sf9 cells were used to amplify the virus. Suspension High Five™ cells were infected with P3 virus for two days at 27°C and 120 rpm shaking. After expression, crude lysates were purified on a HiTrap TALON column (GE Healthcare), cleaved on the column with 3C protease followed by size exclusion chromatography (HiLoad Superdex 200, GE Healthcare). The final product was collected in 25 mM HEPES pH 7.8, 150 mM NaCl, 1 mM TCEP, and 5% glycerol with ~95% purity confirmed by SDS-PAGE. Fractions were checked with SDS-PAGE, pooled, and finally quantified using the NanoDrop spectrophotometer (ND-1000, ThermoFisher Scientific) and the Protein A_280_ program using specific molecular weight and extinction coefficients for either Sox2 or Oct4. Unless otherwise indicated all chemicals were from Sigma-Aldrich.

### Electrophoretic mobility shift assays (EMSAs)

DNA probes were generated by annealing complementary 5’ labeled Cy5 oligos (Metabion International AG) followed by purification from 10% polyacrylamide gels. For binding reactions, WCL (2-4 ug of total protein) or purified proteins were incubated in binding buffer (25 mM HEPES-KOH pH 8, 50 mM NaCl, 0.5 mM EDTA, 0.07% Triton X-100, 4 mg/mL BSA, 7 mM DTT, and 10% glycerol) and 70 nM Cy5-dsDNA at 37°C for 1h. Samples were then loaded onto 6% native polyacrylamide gels (37.5/1 acrylamide/bis-acrylamide) containing 0.3x Tris-borate EDTA and 5% glycerol and run at 10 mA/gel in running buffer of the same composition. Gels were imaged using Fujifilm FLA-9000 fluorescence scanner using (Fujifilm). Fraction bound was determined by densitometry of raw data using Quantity One® (v4.6.7, Bio-Rad) and the following equation for specific bands and then normalized: F_B_ = DNA_bound_/(DNA_bound_ + DNA_unbound_). Half-life was calculated using fraction bound as a function of protein concentration from at least two independent experiments, error bars represent SD.

Equations for decay were determined using nonlinear regression in Prism 7 for Mac (version 7.0a) and only used when goodness of fit, evaluated by R^2^ values, was 0.95 or greater.

For competition experiments, pre-formed protein/DNA or protein/nucleosome complexes (see binding conditions above) were loaded onto native gels (t=O) and then incubated with unlabeled double stranded DNA containing the *Nanog* locus. Protein dissociation was monitored by removing aliquots of the reaction at the given time points and loading them onto a running gel. Protein complex stability was highly variable thus conditions for competition assays were determined empirically and can be found in the tables below.

Supershift assays were run under the same conditions as equilibrium or static EMSAs, see above. After incubation of the proteins with DNA for 1 hour at 37°C, antibody was added to the reactions and incubated at room temperature for 30 min. Antibody/total protein ratios were empirically determined as 1 μg of antibody per 2 μg of total protein. For Oct4 and Sox2 detection, mouse anti-Oct4 from BD Biosciences (611203) and goat anti-Sox2 from Neuromics (GT51098) were used, respectively.

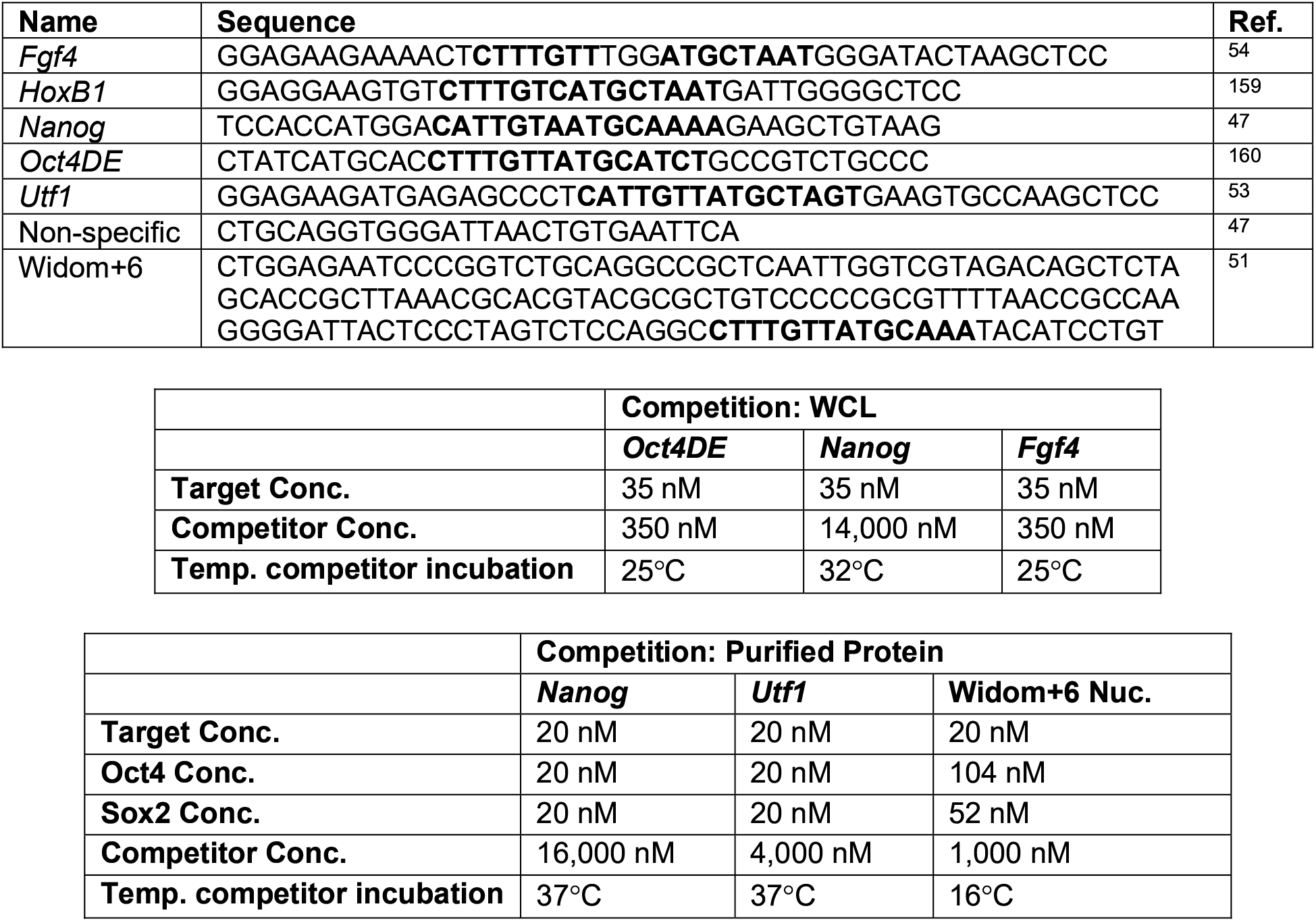

### Nucleosome assembly

The nucleosome DNA sequence Widom +6 consists of 147 bp of the established Widom 601 sequence^161^ with a Sox/Oct motif (**CTTTGTT**ATGCAAAT) at super helical location +6, with the nucleosome dyad being zero (Michael et al 2021). DNA was purchased double stranded from IDT (Coralville) and labeled using Cy-5 conjugated primers via PCR, as previously described^51^. Nucleosomes were assembled from DNA:octamer ratios ranging from 1:1.2-1:1.6 with purified full-length *D. melanogaster* histone octamer^162^ using the salt-gradient dialysis method previously described^163^, final buffer composition: 10 mM HEPES pH 7.6, 50 mM NaCl, 1 mM EDTA, and 0.5 mM DTT. Following dialysis, nucleosomes were heat shifted at 37°C for 2h. Nucleosome quality and concentration were evaluated using native PAGE run with a histone-free DNA standard curve made from the parent DNA. Histone stoichiometry was checked by 22% SDS-PAGE followed by Coomassie staining (R-250; SERVA). Nucleosome were stored at 4°C in the dark and used for no longer than three weeks from the date of assembly.

### Molecular Dynamics Simulation (MDS)

We used the model of the Oct4-Sox2 heterodimer bound to a regulatory DNA element from the Hoxbl enhancer that we previously built^41^ as template for building new models for the Sox2/Oct4, Sox2^A61V^/Oct4, Sox2/Oct6, Sox2^A61V^/Oct6. Using MODELLER (https://salilab.org/modeller/), we adapted the sequences, extended each model of the Oct factor with 4 and 8 residues at the N- and C-termini respectively and similarly, each model of the Sox factor with 4 and 5 residues. We built 100 models for each ternary complex using a “slow” optimization procedure that included a “slow” MD refinement as defined in MODELLER. We ranked the models using an energetic score (“DOPE”) and selected 2 models for each complex for MD simulations. In each of these models, we extended the DNA by 16 and 18 base pairs at the 5’ and 3’ ends using the mouse Hoxbl sequence ^164^. The final sequence in our model was: 5’-AGAGTGATTGAAGTGTCTTTGTCATGCTAATGATTGGGGGGAGATGGAT-3’.

Then, we solvated the systems in a truncated octahedron periodic box of SPCE water with the distance between any protein-DNA atom to the box edges larger than 12 Å. We added 73 neutralizing Na+ ions and 150 mM KCl (225 K+ and 225 Cl- ions). For the ions we used the parameters developed by Li and Merz^165^. We used the Amber-ff14SB^166^ and the Amber-parmbsc1^167^ force fields for proteins and DNA respectively. We energy minimized and equilibrated each system with a protocol described previously^20^. With each model we performed 2 independent, 1.2 μs long MD simulations by assigning different velocity distributions before the equilibration (in total 4 × 1.2 μs = 4.8 μs per system). We applied periodic boundary conditions in the isothermic-isobaric (NPT) ensemble with a timestep of 2 fs. The temperature was maintained at 300 K with Langevin Dynamics (damping coefficient of 0.1 ps-1). The pressure was maintained at 1 atm with the Nose Hoover Langevin Piston method with the period and decay of 1.2 and 1.0 ps respectively. The direct calculation of the non-bonded interactions was truncated at 10 Å and the chemical bonds of hydrogens were kept rigid with the SHAKE algorithm. Long range electrostatics were calculated using the particle mesh Ewald algorithm. All simulations were performed in NAMD^168^. Snapshots were selected for analysis every 10 ps.

The coordination number between two atom selections describes the number contacts between the selections using a continuous switching function with a distance threshold for contact formation as implemented in the COLVAR module of NAMD. The mathematical formula is:

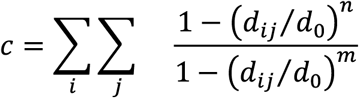

where *i,j* = a pair of atoms, one from each selection; *d_ij_*= the distance between *i* and *j*; *d*_0_= the distance threshold (4.5 Å); *n, m*= exponents describing the switching from contact to no contact (*n* = 6, *m* = 12)

To build the models of the Oct4-Sox2 complexes bound to the Nanog element, we started from the Oct4-Sox2-Hoxb1 model and adapted the sequence using the swapna function in Chimera (https://www.cgl.ucsf.edu/chimera/). The models were minimized and equilibrated with the same procedure. 2 independent simulations, each 2 microsecond long were performed with the same protocol (described above).

### NGS and bioinformatic analysis

For ChlP-seq experiments, Rosa26TA-Gof18 MEFs were infected with titrated volumes of pHAGE2-tetO-Klf4-IRES-Sox2/Sox2^AV^ with or without pHAGE2-tetO-Oct4/Oct6. After 48h, the media was replaced with fibroblast media supplemented with doxycycline (dox). The samples were collected 48h after dox-induction.

RNA-seq and RRBS were performed for human iPSC at passage 10-12, human ESCs (H6^169^ and H9^13^) at passage 35-36 grown in StemFlex media on Matrigel.The sample processing and data analysis for RNA-seq, ChlP-seq, RRBS were performed as described before^22,28,84^.

For RRBS analysis Fastq files were trimmed with Trim Galore (bioinformatics.babraham.ac.uk/projects/trim_galore/) flagging the options --RRBS. The trimmed fastq files were aligned to GRCh38 using bwameth (github.com/brentp/bwa-meth) and methylation metrics were extracted using MethylDackel (https://github.com/dpryan79/MethylDackel), flagging the options --minDepth 10. The genomic coordinates of known imprinted DMRs ^170^ were converted to GRCh38 using the LiftOver tool from UCSC and the methylation levels of CpGs within these regions were extracted with bedtools intersect (bedtools.readthedocs.io/en/latest/content/tools/intersect.html). Finally, the mean methylation level for all CpGs of a given DMR was calculated. The several DMRs corresponding to SNRPN were collapsed as well. Only DMRs which were represented by all samples were taken into consideration for the comparative LOI analysis. A DMR was considered to lose imprinting if it showed less than 30% mean methylation levels.

For footprinting analysis, the publicly available data were aligned to mm10 genome using bowtie2^171^ using “--very-sensitive -X 2000 -no-mixed” options; the mitochondrial and duplicate reads were removed, and the reads were sorted and indexed using samtools^172^; spearman correlation was plotted using deeptools^173^; the peaks were called using macs2 using “-g mm -f BAMPE --call-summits --cutoff-analysis --keep-dup all -B” options^174^; the output of macs2 was used for TOBIAS footprinting analysis^57^ using ENCODE blacklist^175^, JASPAR MEME motif database^176^ with some additional custom motifs^28^.

## Author contributions

SV made the initial discovery, conceived the study, designed and performed most of the experiments, interpreted the results, wrote the manuscript with input from all other authors; HRS advised on the project, raised funding, provided a collaborative environment; CMM established and performed all of the biochemistry work, interpreted the results; VC advised on the project, performed MDS, interpreted the results; GW performed the 4N complementation experiments and interpreted the results; VM performed ChlP-seq experiment and together with RJ analyzed and interpreted the results; TV helped with cloning and reprogramming experiments; GK performed RRBS analysis.

## Acknowledgments

We thank Ingrid Gelker for cell culture support, Claudia Ortmeier for help with RNA preps and western blots, David Obridge and Manuela Haustein for help with teratoma assays, Martin Stehling for FACS, Anika Witten for help with RNA-seq, Novogene for RRBS; Guy Haim from Benvenisty lab for providing the script for e-Karyotyping. We thank Hossein Baharvand and Davood Sabour, Ulrich Martin, Eckhard Wolf and Eva-Maria Jemiller, Heiner Niemann and Monika Nowak-lmialek for sharing human foreskin, cynomolgus, bovine, and porcine fibroblasts, respectively. The work was funded by the Max Planck Society and the Max Planck Society’s White Paper-Project “Animal testing in the Max-Planck-Society”, the ERC (advanced grant proposal 669168), and CiM Pilot Project fund (PP-2017-13). TV was funded by FEBS Summer Fellowship.

## SUPPLEMENTARY INFORMATION

**Fig. S1.**
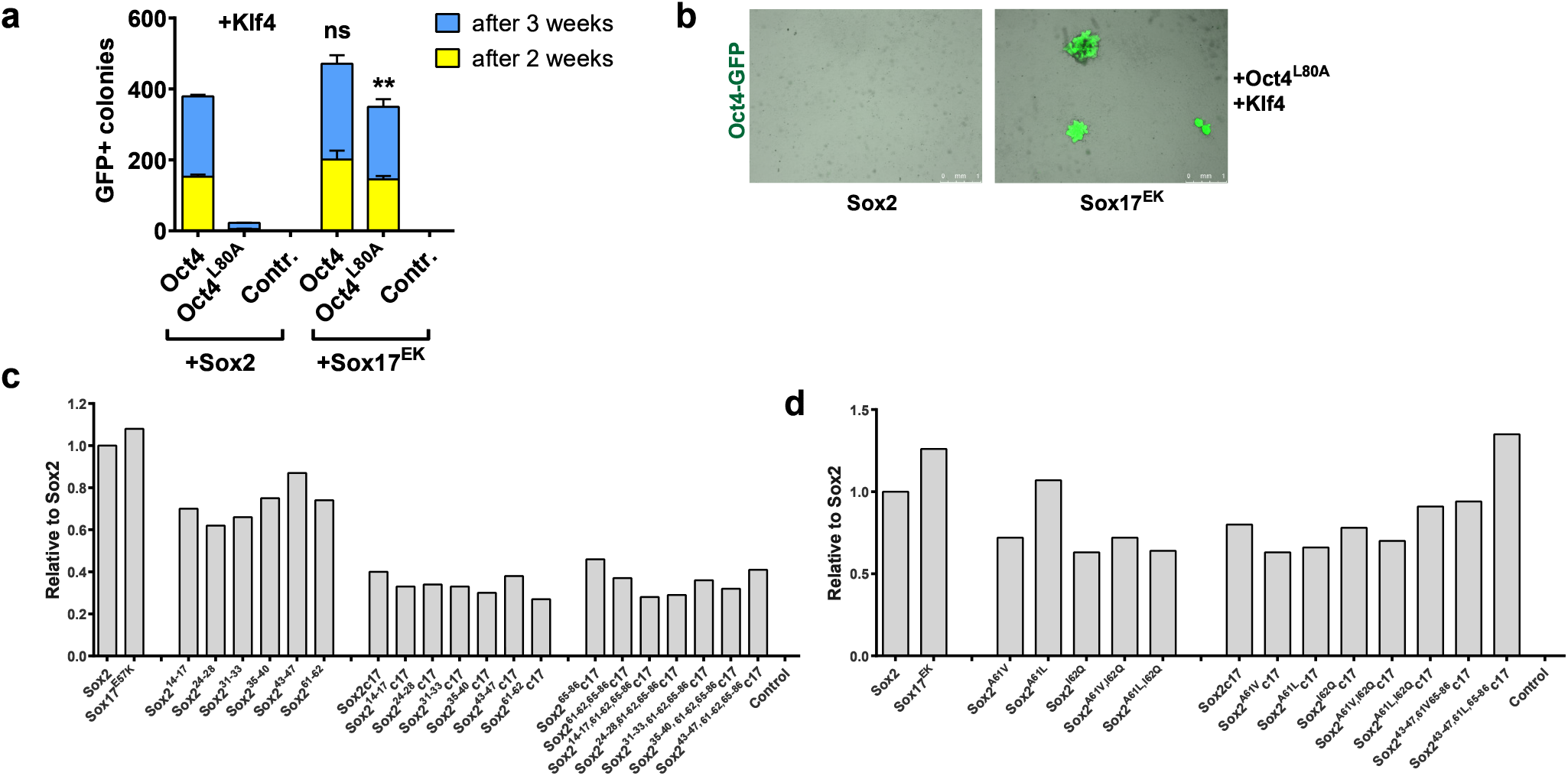
(Related to Fig. 1) **a,** OSK reprogramming of Oct4-GFP reporter MEFs by retroviral monocistronic Sox2 or Sox17^EK^, combined with Klf4 and Oct4^L80A^ linker mutant. Error bars represent SD; n = 3. Statistical significance was calculated with Student’s t-test. b, Representative brightfield and Oct4-GFP merged overview images showing OG2 MEFs reprogrammed with Oct4^L80A^ linker mutant, Klf4, and wild-type Sox2 versus Sox17^E57K^ mutant, 21 dpi, scale=l mm. **c-d**, qPCR titration of the retroviral vectors from Fig. 1d-g.

**Fig. S2.**
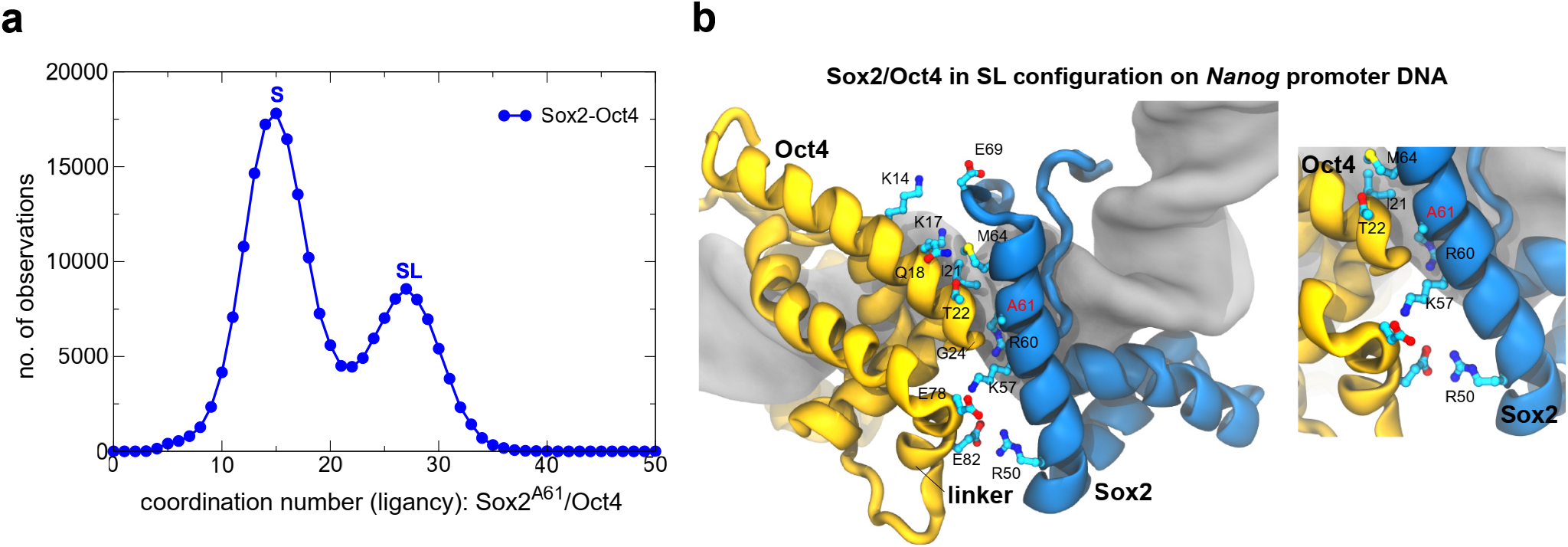
(Related to Fig. 2) **a,** Computer molecular dynamic simulations (MDS) of Sox2/Oct4 heterodimers on *Nanog* promoter DNA. The plots show the coordination numbers (the number of contacts) between the residue A61 in Sox2 (blue) with the entire DNA binding domain of Oct4 molecule. **e**, A model of Sox2/Oct4 binding in POU_S_+Linker (SL) configuration captured from (a).

**Fig. S3.**
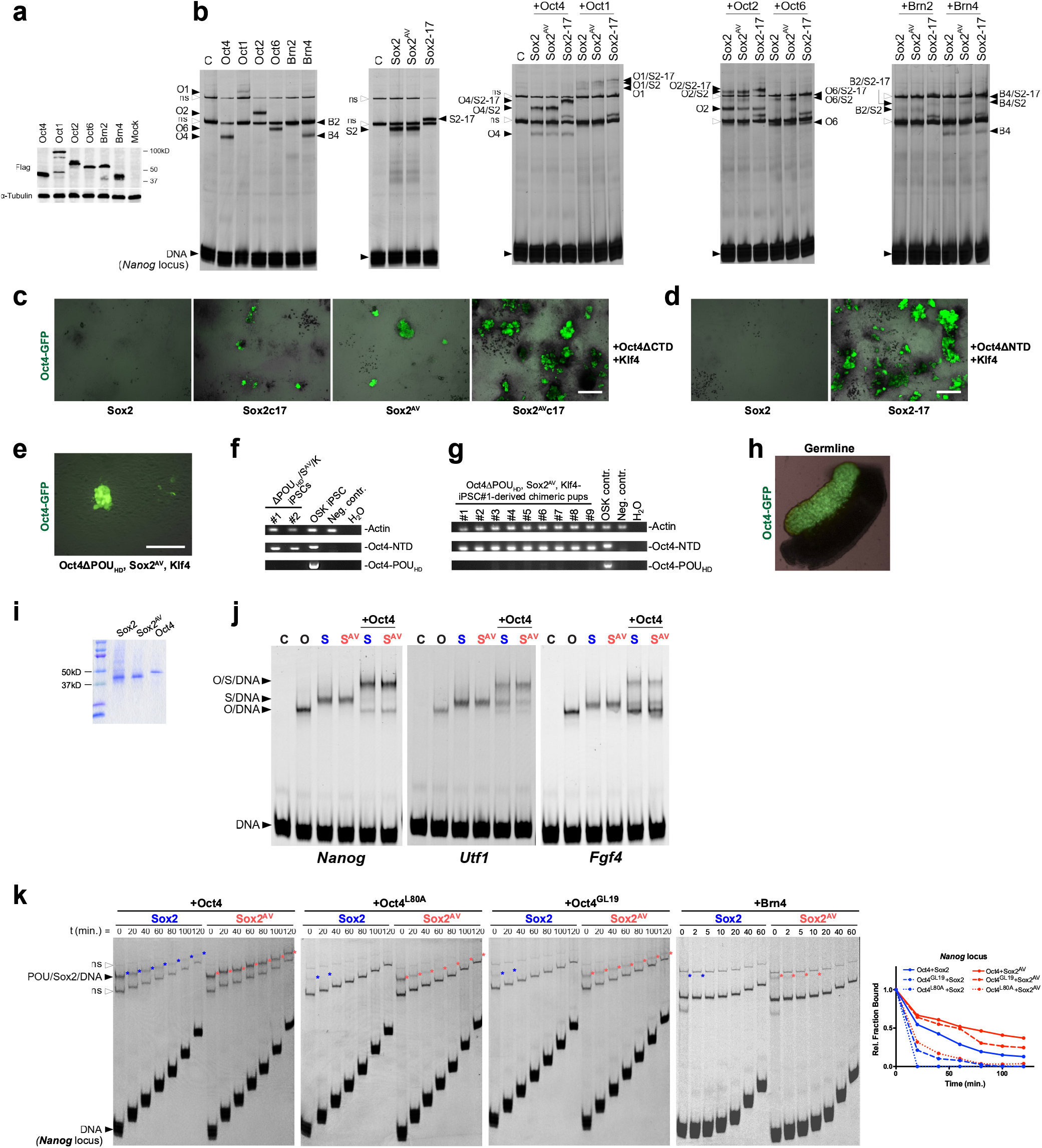
(Related to Fig. 3) **a,** Western blot of whole-cell lysates of HEK293 overexpressing flagged POU factors used in (**b**). **b,** EMSAs of whole-cell lysates from (a) on the *Nanog* promoter locus labeled with Cy5. White arrow heads indicate nonspecific bands (ns) and black arrow heads indicate free DNA or DNA bound by Oct (O/DNA), Sox (S/DNA), or both (O/S/DNA). **c-d**, Representative brightfield and Oct4-GFP merged overview images showing MEFs reprogrammed with Oct4 mutant without the C- (**c**, ΔCTD) or N- (**d**, ΔNTD) terminal transactivator domain, 21 dpi, scale=l mm. **e**, A primary iPSC colony generated by the Oct4 mutant with the POU_HD_ domain removed (except the NLS), scale=l00 μm. **f**, PCR genotyping confirming the identity of two Oct4ΔPOU_HD_/Sox2^AV^/Klf4 iPSC lines, **g**, PCR genotyping of chimeric mice generated by aggregation with Oct4ΔPOU_HD_/Sox2^AV^/K- generated iPSCs. **h**, Brightfield and Oct4-GFP merged image of embryonic day 13.5 gonad dissected from chimeric embryo from (g). **i**, Coomassie stained SDS-polyacrylamide gel of mouse Sox2, Sox2^AV^, and Oct4 from insect cells used in Fig. 3g and Fig. S3 j. **j**, EMSAs of insect cell-purified Sox2 (S, blue), Sox2^AV^ (S^AV^, light red), and wild-type Oct4 on the *Nanog* promoter, *Utf1* and *Fgf4* enhancer DNA elements labeled with Cy5. Arrow heads indicate free DNA or DNA bound by Oct4 (O/DNA), Sox2 (S/DNA), or the heterodimer (O/S/DNA). **k**, Representative kinetic off-rate EMSAs using whole-cell lysates overexpressing full-length Oct4, Oct4^L80A^, Oct4^GL19^, or Brn4 combined Sox2 versus Sox2^AV^ lysates on the *Nanog* promoter locus labeled with Cy5. Following the binding reaction, half-life was determined by adding excess unlabeled *Nanog* element for the indicated time. White arrow heads indicated nonspecific bands (ns) and black arrow heads indicate free DNA or DNA bound by POU/Sox heterodimer.

**Fig. S4.**
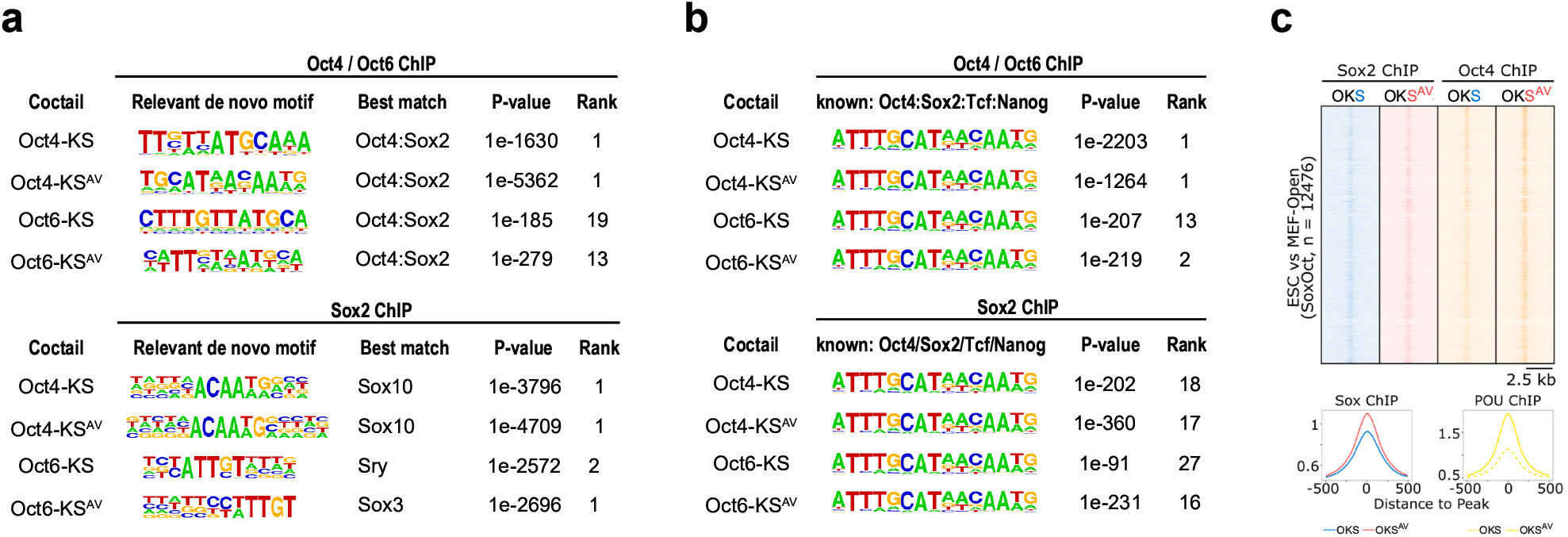
(Related to Fig. 4) **a-b,** HOMER^56^ *de novo* (a) and known *SoxOct* (b) motif analysis showing the enrichment P-value for Oct4, Oct6, Sox2 ChlP-seq. **c**, Sox2 and Oct4 ChlP-seq signal heatmaps for MEFs reprogrammed with tet-inducible OKS at 2 dpi, at the loci containing Sox2/Oct4 footprints in opened chromatin of ESCs versus MEFs determined by TOBIAS footprinting analysis of ATAC-seq data^58^.

**Fig. S5.**
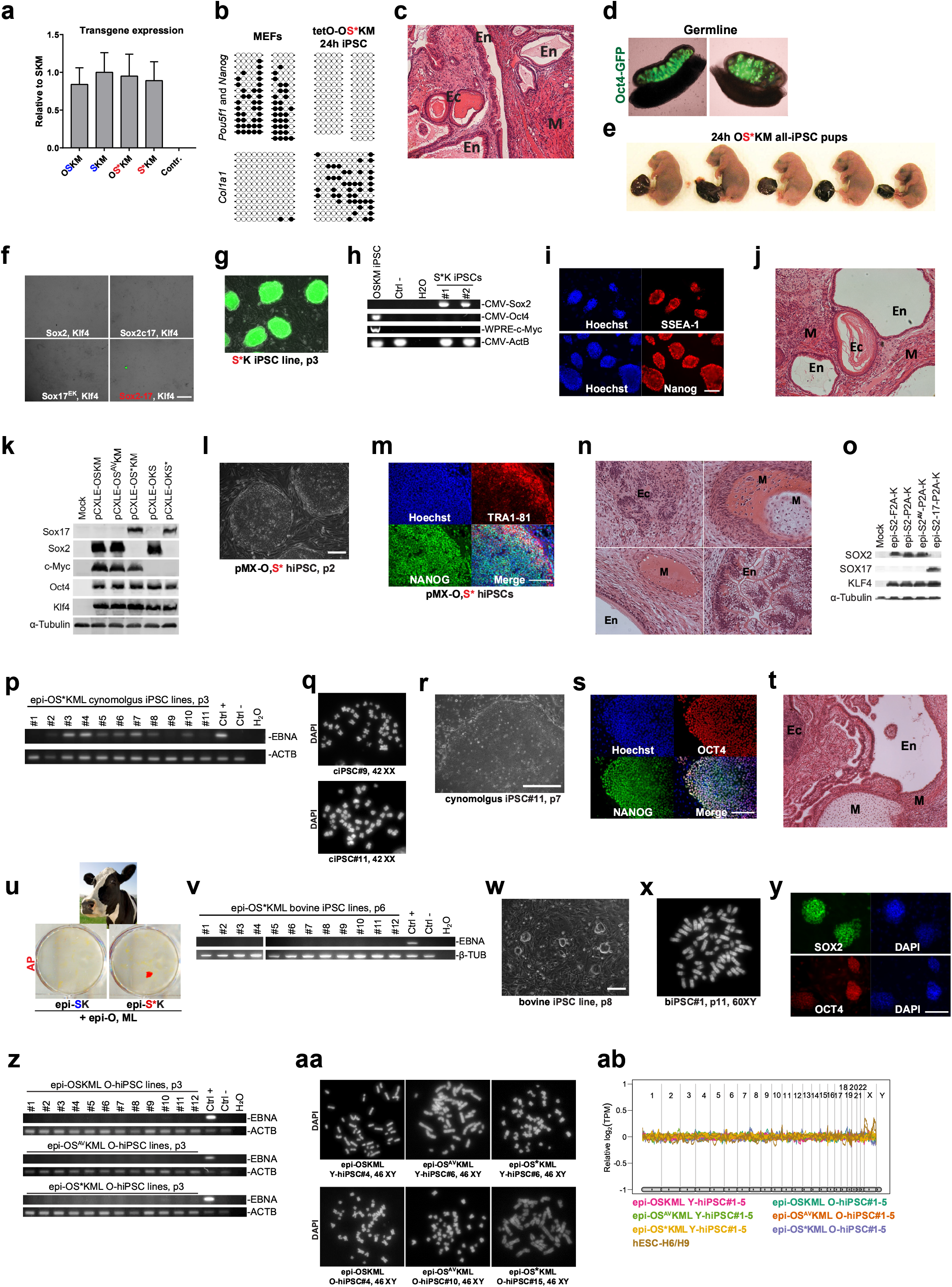
(Related to Fig. 5) **a,** qPCR titration of tet-inducible lentiviral vectors from Fig. 5 b-c after 24h of Dox induction. Error bars represent SD; n = 3. **b**, Bisulfite sequencing analysis of DNA methylation in *Oct4, Nanog*, and *Col1a1* promoters in MEFs and an iPSC line generated by inducing tetO-OS*KM for 24h. **c**, H&E staining of teratoma sections generated with 24h OS*KM iPSCs with representation of the three germ layers (ectoderm - Ec: keratinizing epithelium; mesoderm - M: striated muscles; endoderm - En: cuboidal epithelium). **d**, Bright-field and Oct4-GFP merged images of the gonads from E13.5 24h OS*KM iPSC chimeric embryos. **e**, All-iPSC pups generated by tetraploid (4N) complementation assays with 24h OS*KM iPSC#l line. 12 aggregates were transferred to a pseudopregnant CD-1 (white) female. **f**, Representative brightfield and Oct4-GFP merged overview images showing Oct4-GFP MEFs reprogrammed with tet-inducible lentiviral Sox-T2A-Klf4 vectors carrying wild-type Sox2, Sox2c17, Sox17^EK^, or Sox2-17, 21 dpi, scale<1 mm. **g**, Phase-contrast and Oct4-GFP merged microscopy image of two-factor mouse S*K iPSC clonal line generated in (f) at passage three, scale = 100 μm. **h**, PCR genotyping of two mouse S*K iPSC lines from (f-g). **i**, Immunostaining of a mouse S*K iPSC line for pluripotency markers Nanog and SSEA-1. Nuclei were stained with Hoechst 33342, scale = 100 μm. **j**, H&E staining of teratoma sections generated with S*K mouse iPSC line with representation of three germ layers (ectoderm - Ec: keratinizing epithelium; mesoderm - M: striated and smooth muscles; endoderm - En: cuboidal epithelium). **k**, Western blot of whole-cell lysates from HEK293T transfected with pCXLE-Oct4-P2A-Sox-T2A-KIf4-E2A-cMyc, pCXLE-Oct4-P2A-KIf4-IRES-Sox episomal vectors carrying mouse Sox2, Sox2^AV^, or Sox2-17. **l**, Phase-contrast microscopy image of two-factor human iPSC line generated with monocistronic retroviral OCT4 and SOX2-17 at passage two, scale = 200 μm. **m**, Immunostaining of a human OS* iPSC line for pluripotency markers NANOG and TRA1-81. Nuclei were stained with Hoechst 33342, scale = 100 μm. **n**, H&E staining of teratoma sections generated with OS* human iPSC line with representation of three germ layers (ectoderm - Ec: neural rosettes; mesoderm - M: cartilage, bone, endothelium; endoderm - En: gut and lung epithelium). **o**, Western blot of whole-cell lysates from HEK293T transfected with the original episomal pCXLE-SOX2-F2A-KLF4 construct, as well as generated in this study P2A vectors: pCXLE-SOX2-P2A-KLF4, pCXLE-SOX2^AV^-P2A-KLF4 and pCXLE-SOX2-17-P2A-KLF4. **p,** PCR genotyping of episomal iPSC lines generated from cynomolgus macaque fibroblasts at passage three. **q**, Chromosomal spreads of two integration-free cynomolgus macaque iPSC lines. **r,** Phase-contrast image of integration-free cynomolgus macaque iPSC#11 from panel o at passage seven, scale = 200μm. **s**, Immunostaining of cynomolgus integration-free iPSC line for NANOG and OCT4. Nuclei were stained with Hoechst 33342, scale = 100 μm. **t**, H&E staining of teratoma sections generated with cynomolgus integration-free iPSC line with representation of three germ layers (ectoderm - Ec: neural rosettes; mesoderm - M: cartilage, smooth muscles; endoderm - En: cuboidal epithelium). **u**, A representative whole-well scan of alkaline phosphatase (AP) staining for episomal reprogramming of bovine fetal fibroblasts on day 21 after nucleofection with episomal OSKML (omitting P53 knockdown). **v**, PCR genotyping of episomal OSKML bovine iPSC lines from (u) at passage six. **w**, Phase-contrast image of integration-free bovine iPSC line at passage eight from (v), scale = 200μm. **x**, A representative chromosomal spread of integration-free bovine iPSC line from (v). **y**, Immunostaining of bovine integration-free iPSC line for SOX2 and OCT4. Nuclei were stained with DAPI, scale = 200 μm. **z**, PCR genotyping of episomal iPSC lines generated from dermal fibroblasts of aged male (AG04148) at passage three. **aa-ab,** Karyotyping of human integration-free iPSC lines generated from newborn foreskin fibroblasts (young, Y) or 56-year-old male fibroblast (old, O) using chromosomal spreads (**aa**) or ekaryotyping based on RNA-seq data **(ab).**

**Fig. S6.**
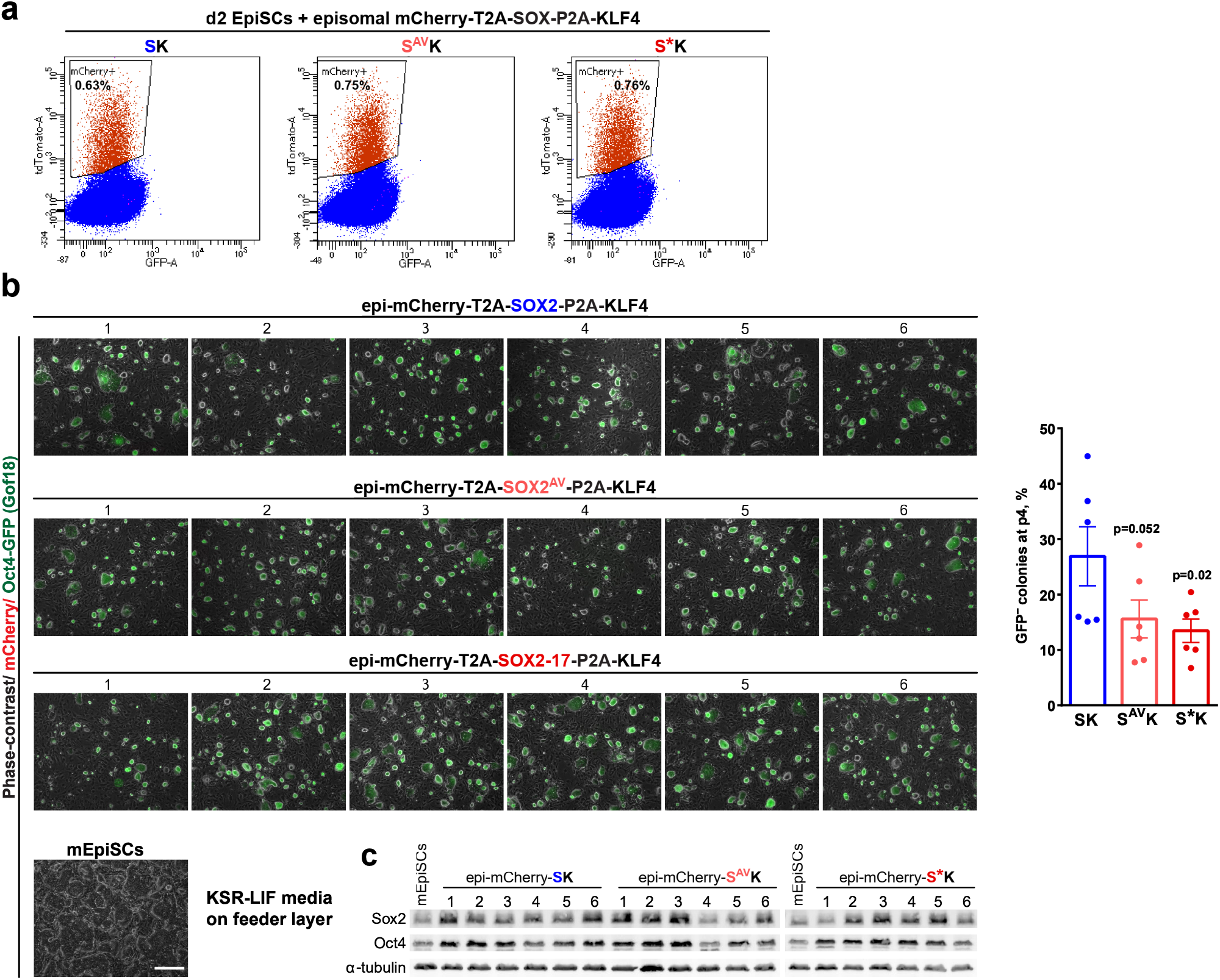
(Related to Fig. 6) **a,** FACS of mEpiSCs transfected with episomal mCherry-T2A-SOX-KLF4 vectors using Lipofectamin Stem reagent at day 2. **b**, Representative phase-contrast/epi-mCherry/Oct4-GFP merged overview images of clonal primed-to-naïve converted mEpiSCs using episomal mCherry-T2A-SOX-P2A-KLF4 vectors grown in KSR-LIF media on a C3H feeder layer, at passage 4, scale=500μm. The same number of cells were plated for each line. At least 100 colonies were quantified from three random overview images of each line, the error bars represent SEM, statistical significance was calculated using Student’s t-test. **c**, Western blot of lysates used in Fig. 6j.

